# Molecular control of replication licensing speed by c-MYC

**DOI:** 10.64898/2026.10.29.762773

**Authors:** Dalia Fleifel, Abid Khan, Abdalla A. Alkhawaja, Arthi Adavikolanu, Sidra Qayyum, Eiman Elkoumy, Jeanette Gowen Cook

## Abstract

A critical prerequisite for DNA replication is genome-wide origin licensing, a process tightly restricted to the G1 cell cycle phase. This restriction means that short G1 cells must license quickly or risk incomplete replication and genome instability, but mechanisms to increase licensing speed are largely unknown. Using quantitative single cell approaches, we identify here a non-canonical role for c-MYC as a potent origin licensing accelerator. Importantly, a point mutation separates c-MYC-induced rapid licensing from the well-known c-MYC gene expression function. c-MYC accelerates licensing through the TRRAP lysine acetyltransferase adapter and GCN5 acetyltransferase. In addition c-MYC has the unexpected ability to induce widespread linker histone H1 eviction. Moreover, directly reducing chromatin-bound histone H1 is sufficient to accelerate licensing. In addition, c-MYC stimulates licensing in heterochromatin, a typically slow-licensing genome compartment. We suggest that global chromatin opening promotes rapid licensing when G1 phase is short such as during early development, rapid progenitor cell proliferation, and adaptation to oncogene activation.

## INTRODUCTION

The eukaryotic cell cycle tightly regulates DNA replication to maintain genome stability. During G1 phase, the Origin Recognition Complex (ORC), CDC6, and CDT1 proteins load MCM2-7 complexes to license replication origins throughout the genome in both the expressed euchromatin and repressed heterochromatin^1,2^. To prevent re-replication and genome instability, licensing is restricted to G1 by the degradation or inactivation of all MCM loading factors once S phase begins^3^. G1 length, the only time available for origin licensing, varies widely among single cells and different cell types^4,5^. Thus, to ensure complete genome duplication, short G1 cells must fully license fast enough to achieve adequate MCM loading before S phase onset, after which no new MCM loading is allowed. An imbalance between the rate of origin licensing and the length of G1 phase can be very detrimental. In particular, slow licensing in a short G1 causes underlicensing, incomplete replication, hypersensitivity to replication stress, and genome instability (Fig. 1A)^6–9^.

**Figure 1.**
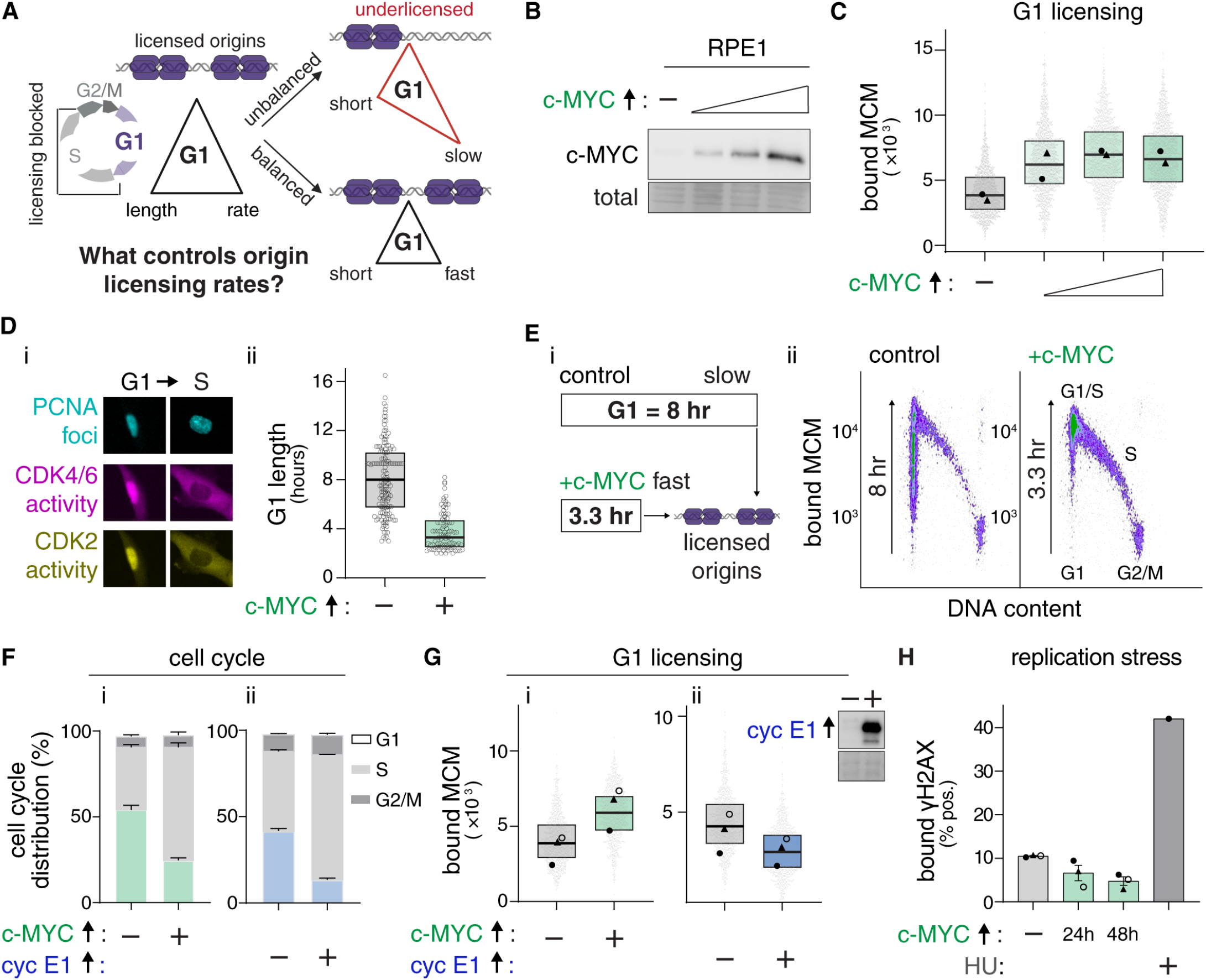
c-MYC accelerates origin licensing. **(A)** Relationships among G1 length, rate of origin licensing (MCM loading), and available origins in S phase. **(B)** Immunoblotting for c-MYC in asynchronously-proliferating RPE1-hTERT cells (2.5, 5, 20 ng/mL doxycycline, 24 hours). **(C)** Chromatin-bound endogenous MCM from G1 phase cells in (B). The aggregate of two biological replicates was randomly downsampled to 3000 cells/sample; line marks the median. Symbols mark the medians of independent replicates. **(D)** i-Representative G1 or S phase cells from live-cell imaging. **ii-G1** lengths in control or c-MYC-expressing cells (control n = 91 cells; +c-MYC n = 153 cells). **(E)** i-Illustration of licensing speed comparisons and the median G1 length from Dii. ii-Analytical flow cytometry for chromatin-bound MCM (“+c-MYC” 20 ng/mL doxycycline, 24 hours); 17,500 cells per plot. **(F)** Cell cycle phase distribution of control or + c-MYC (i: 20 ng/mL) or +cyclin E1 (ii: 100 ng/mL). Data combined from n=3; mean ± SEM. **(G)** Chromatin-bound MCM per cell in G1 from (F), 3000 cells /sample. t-test from three independent biological replicates (not individual cells – see Methods); (i) p=0.004, (ii) p=0.0002. **(H)** Percentage of γH2AX-positive cells; mean ± SEM; t-test from three independent replicates; control vs 24 hours; p=0.15, vs 48 hours; p=0.03.

Despite extensive research into the biochemical process of MCM loading itself, mechanisms controlling the cellular licensing rate are poorly understood. Mammalian pluripotent stem cells have naturally very short G1 phases, yet they also complete genome duplication successfully, in part, because they license origins very rapidly. As pluripotent cells differentiate, licensing slows. Conversely, reprogramming differentiated cells to pluripotency results in rapid licensing^6^. Pluripotent stem cells also have multiple markers of open chromatin, a feature that is associated with both early replication in S phase and early licensing in G1^10,11^. To shed light on the mechanisms that control the rate of origin licensing, we tested the hypothesis that one or more of the original reprograming factors (e.g., OCT4, SOX2, KLF4, and MYC)^12^ can accelerate licensing. In the course of this study, we discovered an unexpected non-canonical ability of c-MYC to induce global histone H1 eviction, and loss of histone H1 accelerates origin licensing.

## RESULTS

### c-MYC accelerates origin licensing

We previously showed that human pluripotent stem cells license origins much faster in their short G1, compared to differentiated counterparts with slower licensing and longer G1^6^. The molecular mechanisms controlling the licensing rate are poorly understood however (Fig. 1A). To quantify MCM loading in G1 in thousands of single cells, we used a previously-established chromatin flow cytometry technique in human retinal pigment epithelial (RPE) cells^6,13^, an untransformed hTERT-immortalized cell line. Briefly, we pulsed asynchronously-proliferating RPEs with EdU to unambiguously distinguish G1 from S phase, then extracted soluble proteins to retain only the chromatin-bound proteins. We detected EdU for DNA synthesis in S phase and stained cells for DNA content (Supplementary Fig. S1A).

To quantify origin licensing, we immunolabelled endogenous MCM2 as a marker of the MCM2-7 complex and quantified chromatin-bound MCM levels in G1 cells (Supplementary Fig. S1B). The loaded MCM complex encircles DNA and has a negligible dissociation rate during G1. Importantly, loaded MCMs are highly stable on chromatin and are not released from DNA until they are actively unloaded in S phase as a downstream consequence of DNA replication termination^14–17^. Thus, MCM loading in G1 is essentially unidirectional rather than a typical combination of DNA binding and dissociation. A G1 population with higher MCM per cell but an unchanged or shorter G1 length indicates faster origin licensing, whereas slower licensing produces a wide distribution of MCM intensity throughout G1^6^. To approximate G1 length, we analyzed cell cycle distributions from the flow cytometry data. The population doubling times did not change, so any decrease or increase in the proportion of G1 cells was the result of changes to G1 length.

To test if one or more of the original reprogramming factors can induce fast licensing, we used a polycistronic construct to inducibly and acutely co-express all four factors, OCT4, KLF4, c-MYC, and SOX2 (Supplementary Fig. S2A)^18^. Expressing all four factors together (OKMS) resulted in substantial G1 phase shortening (Supplementary Fig. S2Bi, C). Simultaneously, most G1 cells expressing all four had high MCM loading despite spending less time in G1 (Supplementary Fig. S2C and Di). However, co-expression of SOX2, KLF4, and OCT4 without c-MYC (OKS) or expressing them individually failed to increase the rate of licensing. (Supplementary Fig. S2Bii, C, Dii and Supplementary Fig. S3A-I). Strikingly, c-MYC expression alone was sufficient to both increase G1 licensing and shorten G1, comparable to the effects of all four factors combined (Supplementary Fig. S2Bi, C, Di versus iii). Even at these modestly increased levels of c-MYC, we observed the same rapid licensing in G1 within 24 hours of induction (Fig. 1B-C), including in an independent clone (Supplementary Fig. S4A-C). Moreover, c-MYC effects were readily detectable within 8 hours of inducing expression and were maximally visible within a single 22-hour RPE cell cycle (Supplementary Fig. S4D-F). The ability of c-MYC to promote G1 progression has been previously reported^5,19,20^, but the effect on origin licensing speed was not previously known.

Comparing licensing speeds requires both MCM loading quantification and G1 length determination. To precisely measure G1 length, we performed live-cell imaging of RPE cells expressing a fluorescent PCNA fusion to mark S phase entry and exit^21^ plus fluorescent CDK2 and CDK4/6 protein kinase activity reporters as measures of G1 progression^22,23^ (Fig. 1Di). Representative film strips from control or c-MYC-overexpressing cells and the imaging workflow are shown in Supplementary Fig. S5A and S5B. Inducible expression of ectopic c-MYC was evident within 2 hours of doxycycline addition (Supplementary Fig. S5Ci), and c-MYC induction in the cell line derivative used for imaging caused fast licensing (Supplementary Fig. S5Cii, D, E). Through time-lapse imaging of asynchronously proliferating cells, we calculated a median G1 length of 3.3 hours in cells expressing c-MYC compared to 8 hours in control G1 cells (Fig. 1Dii). c-MYC-expressing cells spent less than half as much time in G1 as controls, but had loaded just as much MCM by the time S phase started. These results indicate that c-MYC can induce fast licensing (Fig. 1Ei-ii).

Interestingly, while induced cyclin E1 expression shortened G1 similarly to c-MYC, it did not induce fast licensing (Fig. 1F, G)^6,24^. Underlicensing from cyclin E1 overexpression causes DNA replication stress in S phase, an effect we and others have previously shown^6–8,24^. However, the short G1 induced by c-MYC did not increase a marker of replication stress, ɣH2AX, after either 24 or 48 hours of c-MYC induction (Fig. 1H). Although c-MYC stimulated origin licensing in G1, we observed no evidence of re-replication which is a consequence of unscheduled origin licensing outside of G1 phase rather than fast origin licensing (e.g., Fig. 1Eii). We conclude that c-MYC accelerates licensing which avoids acute replication stress in the subsequent S phase that would normally be caused by underlicensing from a short G1.

### Accelerated licensing is independent of licensing factor induction

Because c-MYC is a well-known transcriptional regulator, we tested if its effects on licensing are from directly inducing the genes encoding licensing factors (*MCM2-7, ORC1-6, CDT1,* and *CDC6*) or indirectly such as through stimulating the E2F activator gene expression *(E2F1-E2F3)*. We analyzed mRNA profiles from cells expressing c-MYC for 24 hours versus controls; we analyzed equal numbers of cells to account for possible c-MYC effects on total cellular RNA^25^. Exogenous c-MYC transcript expression in our experiments showed a moderate 3.2-fold increase (Supplementary Table S1). Although some of the candidate MYC-induced gene expression changes were statistically significant (FDR < 0.05), none exhibited a fold-change greater than 1.35 in either direction (Fig. 2A and Supplementary Table S1). In addition, publicly-available transcription profiles comparing chronic c-MYC expression to control conditions in MCF10A cells^26^ showed similar subtle effects on licensing genes (Supplementary Fig. S6 and Supplementary Table S2). Gene set enrichment analysis (GSEA) confirmed that the most upregulated Hallmark gene sets in our data were MYC targets V1 and V2 (Fig. 2B and Supplementary Table S3) as expected^26,27^. Moreover, GSEA of Gene Ontology (GO) Biological Process and Molecular Function terms were enriched for terms related to RNA metabolism and processing and ribosome biogenesis, none of which have known connections to origin licensing dynamics in G1 phase (Supplementary Tables S4, S5). These effects argue against c-MYC accelerating licensing by simply inducing genes for licensing proteins after acute c-MYC expression.

**Figure 2.**
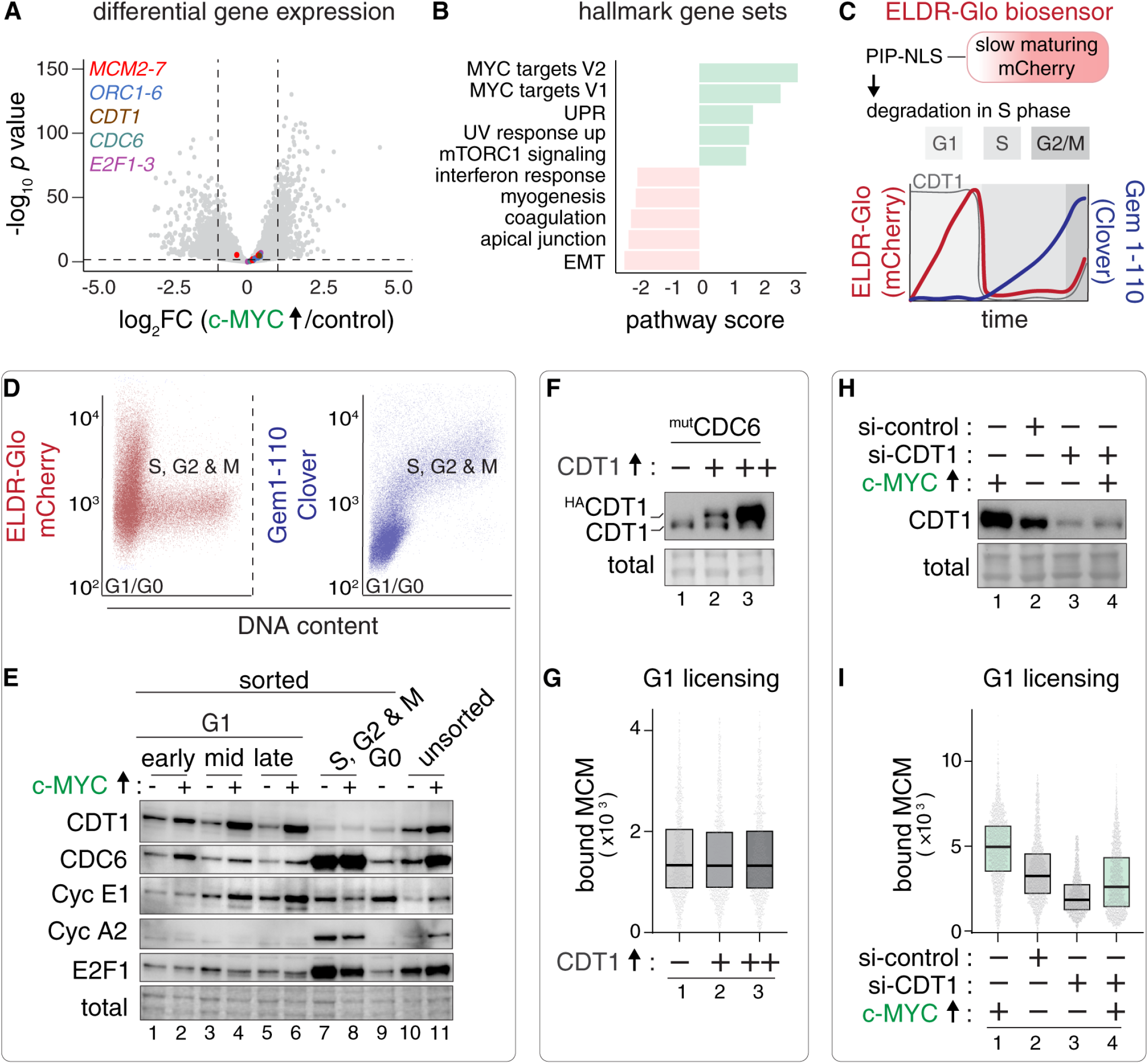
Accelerated licensing is independent of licensing factor induction. **(A)** Differentially-expressed genes for +c-MYC versus control RPE cells (20 ng/mL doxycycline, 24 hours). Licensing factors and activator E2F genes color-coded. Log2(fold change) threshold = ±1, FDR-adjusted p-value threshold = 0.05. Data from 4 independent biological replicates. **(B)** Top five upregulated and downregulated Hallmark pathways by gene set enrichment analysis. UPR: unfolded protein response, EMT: epithelial mesenchymal transition. Pathway score: Normalized Enrichment Score (NES). **(C)** Cell age (ELDR-Glo) and cell cycle (gem 1-110) biosensor dynamics. PIP: PCNA-interacting peptide (degron), NLS: nuclear localization signal. **(D)** Flow cytometry for biosensors in asynchronously-proliferating cells; representative of n>10. **(E)** Endogenous proteins in sorted cells from G1 sub-phases, S, G2 &M, and G0. n=2. **(F)** Endogenous CDT1 or ectopic HACDT1 in +c-MYC RPE1 cells expressing stable CDC6 (mutCDC6) (-) or overexpressing HACDT1 at 15 (+) or 50 ng/mL (++) doxycycline for 24 hours. **(G)** Chromatin-bound MCM per cell in G1 from (F), 1000 cells /sample, n=1. **(H)** Endogenous CDT1 in +c-MYC RPE1 cells treated with control or CDT1 siRNAs for 36 hours, or 20 ng/mL doxycycline for 24 hours as indicated. **(I)** Chromatin-bound MCM per cell in G1 from (H), 3000 cells /sample. n=1.

Because protein changes might not be fully reflected in the mRNA analysis, we also evaluated key licensing proteins. MCM2–7 proteins are abundant, constitutively expressed in proliferating cells, and their chromatin loading rather than abundance is rate-limiting for licensing. For this reason, we did not focus on changes in MCM mRNA or protein levels (Supplementary Fig. S7A). We also analyzed the ORC2, ORC4 and ORC6 subunits as representatives of the ORC complex. Interestingly, although c-MYC increased the licensing rate in G1, it did not increase ORC4 chromatin-bound levels in G1 measured by flow cytometry (Supplementary Fig. S7B). In addition, ORC6 chromatin-bound levels in G1 slightly decreased, rather than increased (Supplementary Fig. S7C). The total levels of ORC2 and ORC4 did not change (Supplementary Fig. S7D).

Licensing is particularly sensitive to the levels of the CDC6 and CDT1 proteins which (unlike MCM2-7) are highly cell cycle-regulated^28,29^, so we focused on the G1 expression of these proteins. To avoid confounding effects from the c-MYC-induced cell cycle changes, we separated asynchronously-proliferating cells by cell cycle phase and G1 subphase using flow cytometric sorting. We utilized a novel “ELDR-Glo” biosensor that reports time in G1 or G0/arrest, (characterized in^30^). We combined it with a second biosensor that only accumulates in S, G2, and early M phases (Fig. 2C)^31^ because it is a degradation substrate of the Anaphase Promoting Complex/Cyclosome (APC/C)^32,33^; the gating scheme and mutually-exclusive biosensor expression are shown in Supplementary Fig. S8A-H. ELDR-Glo is based on a slow-maturing mCherry whose red fluorescence accumulates in G1/G0 as a function of time, and it resets each cell cycle from replication-coupled degradation in S phase (Fig. 2C^30^). With these biosensors we sorted early, middle, and late G1 cells, a subgroup of spontaneously-quiescent cells (G0)^23^, and the combined S through M phase cells (Fig. 2D, E and Supplementary Fig. S8C, E). The sorted populations showed the predicted expression pattern of endogenous cyclin E, cyclin A, and the E2F1 transcription factor (Fig. 2E). Consistent with the much shorter G1 phase, c-MYC expression reduced the maximum G1 ELDR-Glo intensity^19^ (Supplementary Fig. S8E,F). By immunoblotting, c-MYC elevated total CDT1 and CDC6 protein levels (Fig. 2E, compare lanes 1, 3, and 5 to 2, 4, and 6).

Because c-MYC induced the G1 protein levels of CDT1 and CDC6, we tested if this increase explains c-MYC effects by directly co-overexpressing CDT1 and CDC6. We inducibly overexpressed CDT1 in cells harboring a stable version of CDC6 that is resistant to APC^CDH1^-mediated degradation in G1^34^ (Fig. 2F). We had previously shown that stable CDC6 slightly increases the rate of licensing^6^. Although c-MYC induced CDT1 protein in G1 cells (Fig. 2E), even supraphysiological CDT1 levels combined with stable CDC6 did not increase the rate of licensing in RPE cells (Fig. 2G)^6^. Furthermore, partial CDT1 depletion prevented c-MYC from inducing CDT1 (Fig. 2H compare 2 and 3) and reduced MCM loading in control cells as expected (Fig. 2I compare samples 2 and 3), but c-MYC still induced G1 licensing (Fig. 2I, compare 3 and 4). From these results, we conclude that the potent licensing acceleration by c-MYC was not simply a consequence of increasing CDC6 and CDT1.

### A mutation separating c-MYC-induced gene expression from licensing acceleration

To dissect the mechanism by which c-MYC increases licensing speed, we generated multiple inducible mutant c-MYC cell lines. The c-MYC protein includes an N-terminal transcriptional activation domain (NTD) and a C-terminal DNA-binding domain (CTD) (Fig. 3A)^35,36^. Deleting either the NTD or CTD abolished the effects of c-MYC on licensing and cell cycle phase distribution (Supplementary Fig. S9 A-C). Within the NTD, the conserved “MYC Box zero” (MB0) domain has been implicated in transcriptional regulation through various protein-protein interactions such as TFIIF^27^. In contrast to deleting the entire NTD, c-MYC lacking only the 18-24 amino acid MB0 was nearly as capable as wild-type (WT) c-MYC for increasing licensing speed (Supplementary Fig. S9 D-F). Therefore, MB0-mediated interactions are not required for the c-MYC licensing function. The licensing phenotype of this mutant additionally argues against a purely transcriptional mechanism.

**Figure 3.**
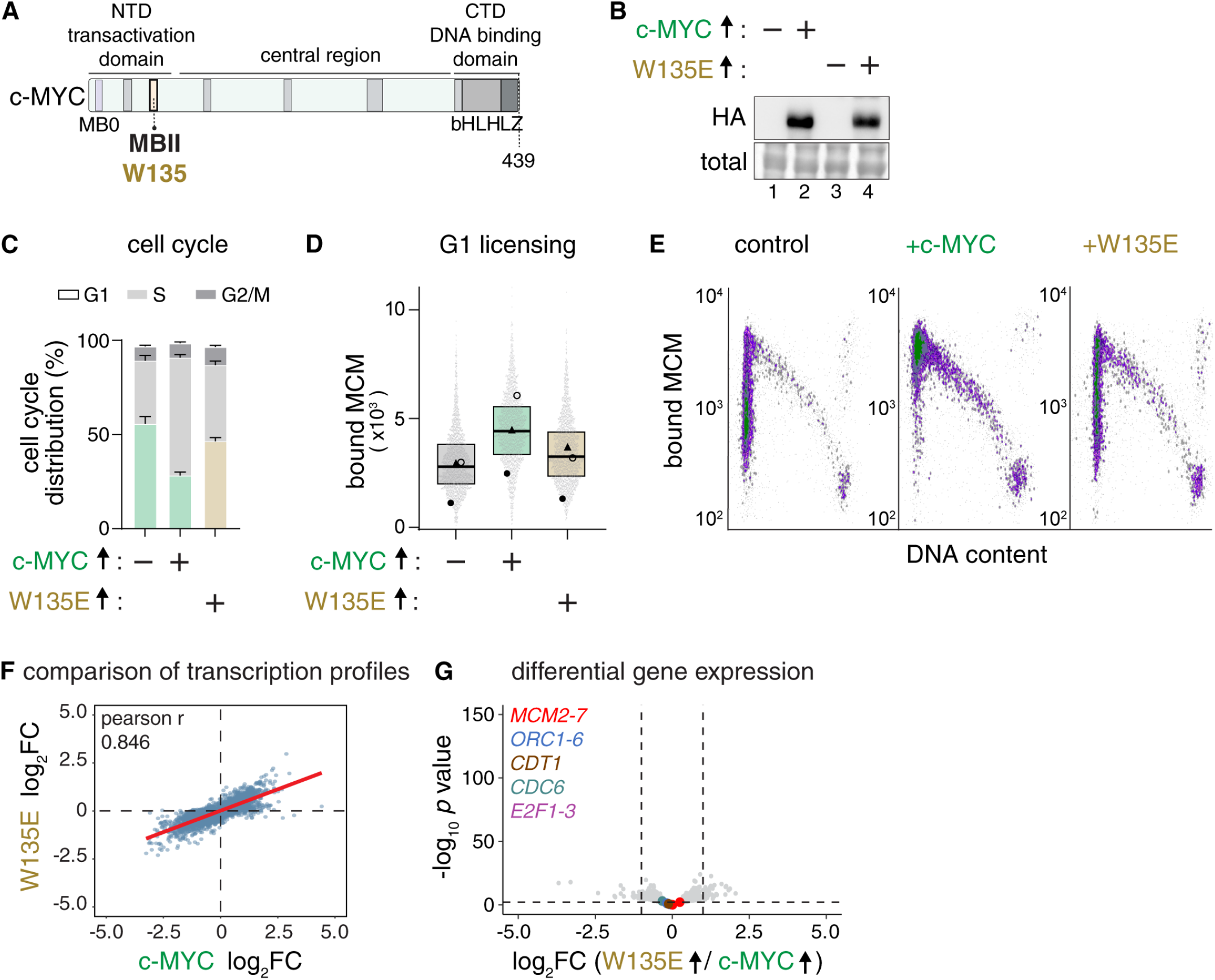
A mutation separating c-MYC-induced gene expression from licensing acceleration. **(A)** Human c-MYC domain structure marking the N-terminal domain (NTD), central region, and C-terminal domains (CTD) as horizontal lines, and MYC-Boxes as vertical bars; bHLH-LZ is basic helix-loop-helix leucine zipper. **(B)** Immunoblotting for HA to detect inducible 2xHAc-MYC or 2xHAc-MYC harboring a single point mutation at tryptophan 135 (2xHA c-MYC W135E) in control or doxycycline-treated RPE1 cells (20 ng/mL, 18-24 hours). Representative of n > 3. **(C)** Cell cycle phase distribution from the same conditions in (B). Data are combined from n=3; mean ± SEM. **(D)** Chromatin-bound MCM per cell in G1 from (C), 4,500 cells /sample. Average change in median MCM intensity relative to control of n=3: WT c-MYC = 1.91; W135E = 1.44. **(E)** Flow cytometry from replicate 1 in (D) for control, WT c-MYC, c-MYC W135E (20 ng/mL, 18 hours). 9000 cells/plot; n>3. **(F)** Pearson correlation (r=0.846) between W135E and WT c-MYC log2(fold change) for 7704 genes with FDR-adjusted p < 0.05 in either condition. **(G)** Differentially-expressed genes for W135E versus WT c-MYC (20 ng/mL, 24 hours) as in Fig. 2A. Log2(fold change) threshold = ±1, FDR-adjusted p-value threshold = 0.05. Data from 4 biological replicates.

Within the NTD, the MBII (MYC Box 2) domain plays a crucial role in MYC-mediated chromatin modifications, particularly through interaction with lysine acetyltransferases^27^. Mutations of the conserved tryptophan W135 in MBII disrupt c-MYC interaction with various co-regulators and also inhibit MYC-induced oncogenic transformation^37,38^. Strikingly, expressing the W135E c-MYC mutant to levels comparable to WT c-MYC had only modest effects on both origin licensing in G1 and G1 length (Fig. 3C-E). Moreover, W135E c-MYC is licensing-impaired relative to WT c-MYC despite similar expression of individual licensing genes (Supplementary Fig. S10A) and induction of G1 CDT1 protein levels (Supplementary Fig. S10B, C).

For a comprehensive comparison of WT and W135E c-MYC, we analyzed mRNA profiles of cells expressing similar levels of WT c-MYC and W135E. Analyzing the union of differentially expressed genes (FDR < 0.05 in either condition) revealed a strong positive correlation of log2fold changes (Pearson r=0.843), indicating highly similar transcriptional responses (Fig. 3F and Supplementary Tables S1, S6, S7). Only 61 genes exhibited a fold change greater than or equal to 2 in either direction between the two conditions, and none were genes for core licensing proteins (Fig. 3G and Supplementary Table S7). Analysis of publicly-available transcription profiles comparing chronic W135E c-MYC versus WT expression in MCF10A cells indicates that both have comparably subtle effects on licensing and E2F upstream activator genes (Supplementary Fig. S10D and Supplementary Table S8)^26^. In addition, analysis of our dataset comparing W135E c-MYC versus WT showed that the differentially-enriched pathways or genes were not obviously related to origin licensing (Supplementary Tables S9-S11). Taken together, our results suggest that the mechanism by which c-MYC accelerates licensing is not through its ability to change mRNA expression. The W135E mutation in MBII separates the well-known gene expression functions of c-MYC from this unanticipated role in origin licensing speed control.

### Licensing acceleration by c-MYC depends on TRRAP and GCN5

Previous studies that focused on W135 in other settings provided clues to possible mechanisms for c-MYC-induced licensing^37,38^. Mutations to W135 impair c-MYC interaction with the TRRAP adapter protein (Fig. 4A)^26,39^. To test if TRRAP is involved in c-MYC-stimulated rapid licensing, we depleted TRRAP using siRNA. Strikingly, TRRAP depletion clearly reduced the ability of c-MYC to induce fast licensing, though it did not prevent c-MYC from inducing a shorter G1 (Fig. 4B, C). Thus, the rate of licensing is not increased because c-MYC shortens G1, and TRRAP is important for c-MYC to increase licensing speed.

**Figure 4.**
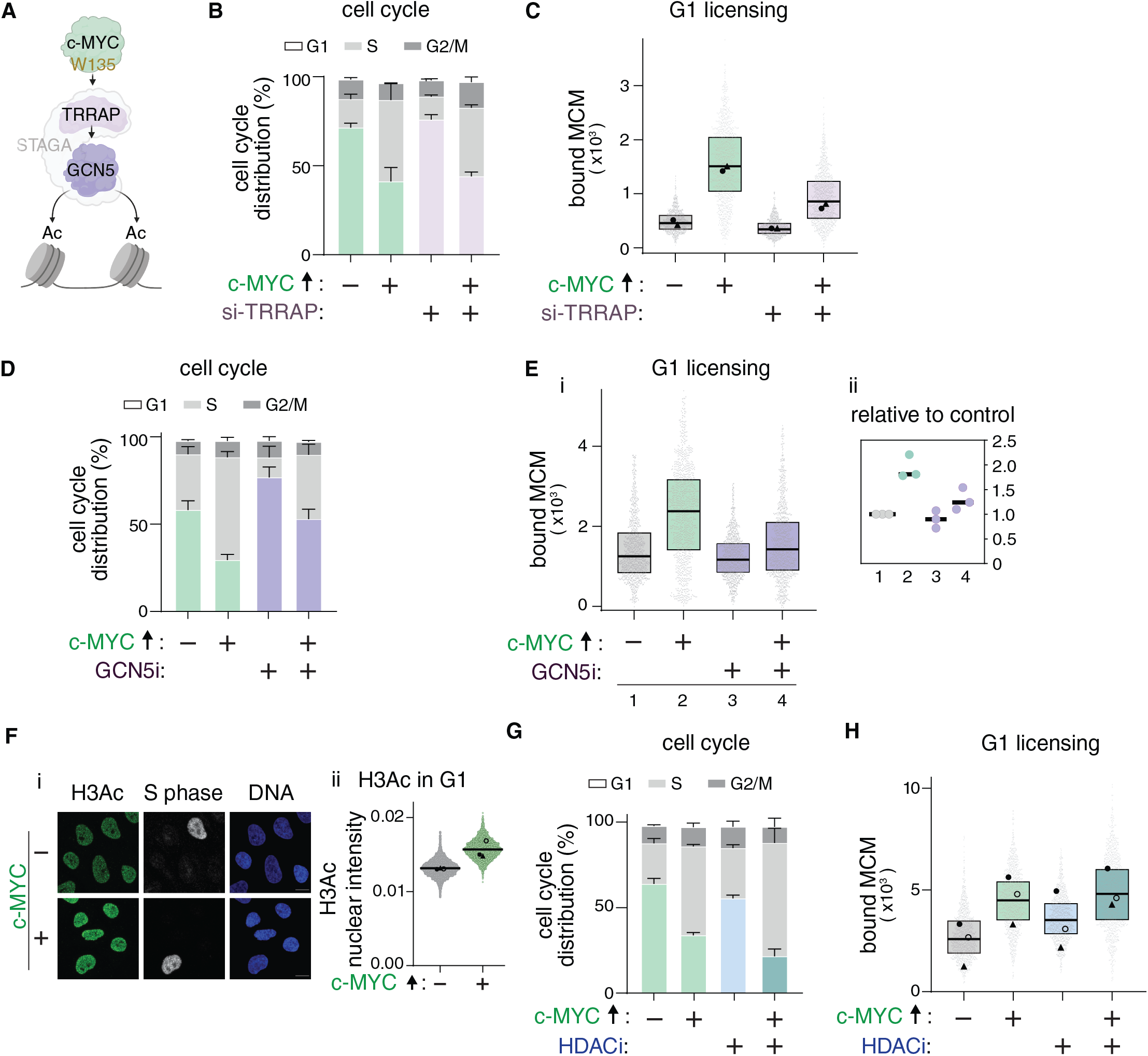
Licensing acceleration by c-MYC depends on TRRAP and GCNS. (A) Illustration of relevant interactions. (B) Cell cycle phase distributions; mean± SEM. +c-MYC RPE cells were treated with control or TRRAP siRNAs for 48 hours, then 20 ng/ml doxycycline for the final 19-24 hours as indicated, n=2. (C) Chromatin-bound MCM per cell in G1 from (B), 2000 cells /sample, n=2. (D) Cell cycle phase distributions; mean± SEM. +c-MYC RPE cells treated with GCN5 inhibitor (MB-3) at 200 µM or 20 ng/ml doxycycline for 16-24 hours as indicated, n=3. (E) i-Chromatin-bound MCM per cell in G1, representative from (D), 1000 cells /sample. t-test; p=0.01. ii-Normalized fold change relative to control. (F) i-lmmunofluorescence for pan-H3 acetylation (20 ng/ml, 20 hours). EdU marks S phase. Scale bar 10 µm, n=3. ii-Quantification of G1 cells in (i), control: 9822, + c-MYC: 7656, n=3. t-test; p=0.057. (G) Cell cycle phase distributions; mean± SEM. +c-MYC RPE cells were treated with the HDACi (TSA) for 6 hours at 300 nM, washed, then 20 ng/ml doxycycline was added for 16 hours, n=3. (H) Chromatin-bound MCM per cell in G1 from (G), 3000 cells /sample, n=3. t-test for control vs HDACi, p=0.09; HDACi vs HDACi + c-MYC, p=0.09.

Two related MYC family members, L-MYC and N-MYC have domain structures that are similar to c-MYC^36^ (Supplementary Fig. S11A). To determine if they also have licensing functions, we inducibly expressed either L-MYC or N-MYC in RPE cells (Supplementary Fig. S11 B and E.) Although N-MYC could shorten G1 and increase the rate of licensing like c-MYC, L-MYC did neither (Supplementary Fig. S11C-I). L-MYC has an MBII domain that is similar to both c-MYC and N-MYC, however a previous study found that L-MYC only weakly interacts with TRRAP, compared to N-MYC or c-MYC^40^. This distinction further supports a role for TRRAP in c-MYC-mediated licensing acceleration.

TRRAP recruits multiple lysine acetyltransferases that can modify histones to stimulate gene expression^41,42^ (Fig. 4A). Given strong links between chromatin compartments (e.g. active vs inactive) and both origin licensing and origin firing^8,43–45^, we considered that c-MYC may induce rapid licensing by pervasively altering chromatin. TRRAP interacts with the GCN5 and NuA4/TIP60 acetyltransferases, and TRRAP interaction with c-MYC brings these enzymes to chromatin^41,42,46^. We tested if either of these enzymes is required for c-MYC to increase licensing using pharmacological inhibitors. GCN5 inhibition consistently diminished c-MYC-induced licensing acceleration and G1 shortening (Fig. 4D, E), whereas TIP60 inhibition did not (Supplementary Fig. S12A, B). We next investigated the involvement of alternative, TRRAP-independent HATs by inhibiting HBO1 and p300. Intriguingly, while HBO1 was dispensable for c-MYC-mediated G1 shortening and fast licensing, p300 inhibition markedly blunted the licensing effect (but not the cell cycle effect, Supplementary Fig. S12C-F), suggesting a parallel requirement for p300.

Multiple previous gene expression and cellular transformation studies showed that c-MYC induces histone hyperacetylation^27,41,47,48^. Hyperacetylation disrupts the electrostatic interactions between histones and DNA, resulting in a less compact chromatin structure^49,50^ which might facilitate rapid origin licensing. The contributions from at least two enzymes known to acetylate histones prompted us to examine histone acetylation in the context of c-MYC-induced licensing acceleration. Because all genomic regions must be licensed for genome duplication, and our licensing assay quantifies total MCM per cell rather than at specific loci, we hypothesized that licensing-related c-MYC effects on G1 chromatin should be widespread. We first analyzed G1 cells using an antibody that recognizes multiple acetylation sites on histone H3 (pan-H3ac) and quantified the mean fluorescence intensity of histone H3Ac by immunofluorescence. Consistent with previous reports^27,48^, c-MYC induced pan-H3Ac, including in G1 nuclei (Fig. 4Fi-ii). Because GCN5 acetylates histone H3 at lysines 9 and 14 (H3K9ac and H3K14ac)^51,52^, we also tested global histone H3K9 and H3K14 hyperacetylation using flow cytometry. We treated with the histone deacetylase inhibitor (HDACi), trichostatin A (TSA), as a positive control for detecting strongly increased histone acetylation by this method (Supplementary Fig. 13A, B). We found that c-MYC modestly induced both histone H3K14 and H3K9 acetylation (Supplementary Fig. S13C, D).

To test if increased hyperacetylation can directly affect licensing speed, we treated control cells or c-MYC overexpressing cells with the TSA histone deacetylase inhibitor. We treated cells with HDACi for just 6 hours to avoid its long-term toxic effects. Strong hyperacetylation by HDACi treatment alone slightly accelerated licensing, but not by as much as c-MYC expression (Fig. 4G, H). We note that HDACi also induced markedly more histone acetylation in less time than c-MYC did (Supplementary Fig. S13A, B), but HDACi was not as potent as c-MYC for increasing origin licensing speed. Moreover, adding elevated c-MYC expression to histone hyperacetylation increased licensing more than HDACi alone, but hyperacetylation by HDACi did not augment c-MYC-induced licensing (Fig. 4H).

Surprisingly, the licensing-impaired W135E c-MYC mutant also induced global pan-histone H3 acetylation comparable to WT c-MYC (Supplementary Fig. S14A), yet W135E was not as effective as WT c-MYC for increasing licensing speed (Figure 3). In addition, W135E expression induced licensing to approximately the same degree as histone deacetylase inhibitor treatment (Supplementary Fig. S14B, samples 2 and 3). Importantly combining HDACi treatment with W135E overexpression did not recapitulate the effects of WT c-MYC (Supplementary Fig. S14B, samples 2 versus 4) which we interpret as W135E and the inhibitor affecting licensing speed by similar mechanisms. W135E is less potent than WT c-MYC for licensing acceleration although the mutant induces histone acetylation effectively. Taken together, these results indicate that hyperacetylation cannot fully explain how WT c-MYC increases the speed of origin licensing.

### c-MYC induces linker H1 eviction which is itself sufficient to accelerate licensing

When considering additional mechanisms for c-MYC to promote rapid licensing, we recalled that fast-licensing pluripotent stem cells have open chromatin that is not only hyperacetylated but also has very low chromatin-bound linker histone H1^11,53^. Linker histone H1 contributes to higher-order chromatin structure and chromatin compaction^54^, thus low histone H1 might support rapid licensing through global chromatin opening. To test this idea, we identified which of the multiple human histone H1 genes are relevant based on their expression; our mRNA profile analysis indicated that histone H1X is the most abundant histone H1 variant in RPEs (Supplementary Fig. S15A). We then examined chromatin binding of the dominant histone H1 variant in response to c-MYC expression.

Strikingly, elevated c-MYC expression significantly decreased histone H1X on chromatin in G1 cells (Fig. 5A, B). Moreover, the licensing-impaired W135E mutant had no effect on histone H1X chromatin binding (Fig. 5B). Furthermore, c-MYC reduced the overall histone H1X signal on chromatin throughout the nucleus (Fig. 5C). Most importantly, direct histone H1X depletion by siRNA, *without* c-MYC overexpression, was itself sufficient to increase the rate of MCM loading (Fig. 5D-F), strongly suggesting that the physical loss of histone H1X from chromatin is enough to increase origin licensing speed. Because depleting the major histone H1 variant in RPE cells accelerated licensing (Fig. 5E), we consider histone H1 eviction to be a major component of the mechanism of c-MYC-induced rapid origin licensing.

**Figure 5.**
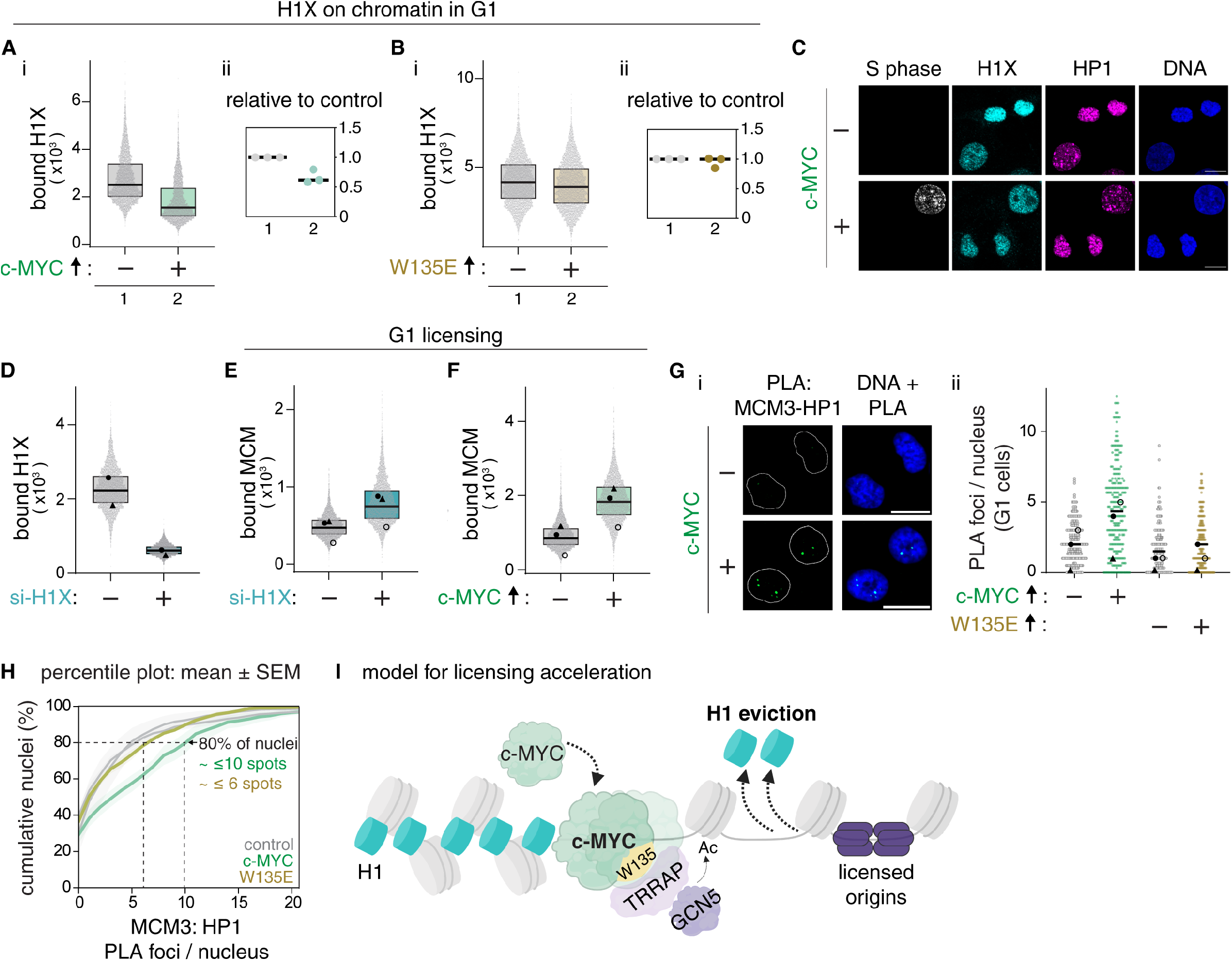
c-MYC induces linker H1 eviction and accelerates licensing in heterochromatin. **(A)** i-Chromatin-bound endogenous H1X levels in G1 (20 ng/mL, 24 hours), 5,000 cells/sample, representative of n=3. t-test of median intensities; p=0.048. ii-Normalized fold change. **(B)** As in (A) for c-MYC W135E; p=0.43. **(C)** Chromatin-bound H1X and HP1 in +c-MYC RPE (20 ng/mL, 20 hours). EdU marks S phase; scale bar, 10 µm. Representative of n=3. **(D)** Flow cytometry for chromatin-bound H1X in G1 after 48 hours histone H1X siRNA, 10,000 cells/sample, n=2. **(E)** Chromatin-bound MCM per cell in G1 from conditions in (D), 14,000 cells/sample, n=3. t-test; p=0.0058. Average 1.64 fold-change. **(F)** Chromatin-bound MCM per cell in G1 from conditions in (A), 14,000 cells/sample, n=3. t-test; p=0.028. Average 2.28 fold-change. **(G)** i-Representative proximity ligation assays (PLA) for chromatin-bound MCM3 and HP1 in +c-MYC cells (20 ng/mL, 24 hours). Scale bar, 10 µm, n=3. ii-MCM3-HP1 PLA foci/ G1 nucleus, n=3: total foci from control=659, WT c-MYC=619, control=528, W135E=525. Lines mark medians. **(H)** Replicate-aware percentile empirical cumulative distribution function (ECDF) for the cumulative percent of nuclei vs PLA foci/nucleus for each condition in (Gii). Shaded areas are SEM of n=3. t-test for median foci/nucleus; control vs WT; p=0.006, control vs W135E; p=1. **(I)** Model for origin licensing acceleration.

To test these findings in another cell line, we engineered the non-transformed human pancreatic nestin-expressing cell line (HPNE) to inducibly express HA-tagged c-MYC. Notably, HPNE cells overexpressing c-MYC also had accelerated licensing (Supplementary Fig. S16A, B). We examined chromatin-bound endogenous histone H1 by chromatin fractionation and immunoblotting in these HPNE cells as a complement to the flow cytometry method. We used either a pan-H1 antibody that recognizes multiple variants or antibody to histone H1.0 which is the most abundant histone H1 variant in HPNE cells based on publicly-available RNA profiles^55^. Similar to RPE cells, chromatin-bound histone H1.0 in c-MYC-expressing HPNE cells was substantially lower than in control cells (Supplementary Fig. S16C). We also tested if drug-induced hyperacetylation promotes histone H1 eviction (Supplementary Fig. S17A) or if RNAi-induced histone H1 depletion affects H3K14 acetylation (Supplementary Fig. S17B). Interestingly, neither of these manipulations suggested that these two chromatin features regulate each other under our conditions, indicating that c-MYC affects chromatin structure by two independent mechanisms.

### c-MYC accelerates licensing in heterochromatin

We observed that histone H1X was uniformly localized in nuclei regardless of cell cycle phase, including in regions enriched for the heterochromatin marker HP1 (Fig. 5C and Supplementary Fig. S15B). Our assays quantified substantial increases in total chromatin-bound MCM to license origins and overall reductions in chromatin-bound histone H1. For this reason, we assume that the chromatin changes are widespread and include both the active and inactive chromatin compartments. We previously demonstrated that HP1-marked heterochromatin is particularly vulnerable to the consequences of underlicensing because heterochromatin is licensed relatively slowly in G1^8,56^. We thus hypothesized that c-MYC-mediated licensing acceleration includes effects in heterochromatin.

To test this idea, we performed proximity ligation assays (PLA) of endogenous chromatin-bound MCM3 and heterochromatin protein HP1. This method detects spatial associations between chromatin-bound MCM3 and HP1 as a readout for heterochromatin licensing. We quantified PLA foci in G1 RPE cells inducibly expressing either WT c-MYC or the licensing-impaired W135E mutant. Strikingly, we observed a significant increase in the number of MCM3-HP1 PLA foci in G1 cells expressing WT c-MYC compared to control G1 cells; the W135E mutant was entirely unable to stimulate MCM3-HP1 proximity (Fig. 5Gi, ii). To assess the distribution of PLA foci in the population, we utilized empirical cumulative distribution functions. This analysis revealed a uniform rightward shift toward more MCM3-HP1 foci per cell in WT c-MYC-expressing cells relative to both control and W135E-expressing cells, indicating an increase in cells with more MCM3-HP1 foci. Consistent with this shift, cells with fewer foci were enriched in control or W135E cells, relative to WT c-MYC cells. For example, 80 percent of control or W135E-expressing cells had fewer than or equal to six MCM3:HP1 PLA foci, whereas the same fraction, 80 percent, of WT c-MYC-expressing cells had more than or equal to ten foci (Fig. 5H). Considered another way, the 20% of control or W135E cells with the most foci in the population only had more than six foci whereas the 20% of the WT c-MYC cells with the most foci had more than ten.

Interestingly, we also detected similar close proximity to HP1 of both WT and W135E c-MYC (Supplementary Fig. S18A-C). Thus, the weak ability of W135E to promote licensing in heterochromatin cannot be attributed to a defect in MYC interaction with heterochromatin. Collectively, our data support a model wherein elevated levels of c-MYC interact with chromatin broadly, including in heterochromatin to drive rapid origin licensing through two independent mechanisms: TRRAP-GCN5-dependent hyperacetylation and linker H1 eviction (Fig. 5I).

## DISCUSSION

Previous studies have shown DNA replication effects from c-MYC manipulation, but these studies focused on the ability of MYC to stimulate the S phase events of origin initiation, and not the prerequisite origin licensing process in G1^57–59^, or they examined licensing at a single origin^60^, but not licensing dynamics. The general correlation of some features of euchromatin with origin activity is established^8,43,61,62^, but a licensing role for linker histone H1 was not previously known, particularly in mammalian cells or with respect to the speed of licensing.

Multiple observations from our study and others argue against c-MYC accelerating licensing through its most well-known role as a transcriptional regulator: **(1)** c-MYC induced minimal changes in mRNAs encoding licensing proteins or the E2F activators; **(2)** although c-MYC is detectable at the *CDT1* promoter^63^ and it can induce CDT1 and CDC6 protein levels in G1, overexpressing CDT1 does not induce fast licensing even combined with CDC6; **(3)** the MB0 domain is required for many transcriptional changes induced by c-MYC^27^, yet is dispensable for fast licensing; **(4)** the W135E mutant is severely compromised for licensing function, yet it binds TATA binding protein (TBP)^64^, and its acute transcriptional effects are remarkably similar to WT c-MYC (shown here and also^37^); the few genes uniquely sensitive to MBII perturbations are primarily involved in ribosome biogenesis and RNA processing/metabolism (Supplementary Table S10 and ^27^); **(5)** chromatin-based mechanisms of c-MYC induced gene expression include histone acetylation^47^, yet directly inducing histone hyperacetylation only modestly accelerated licensing; **(6)** gene expression effects from depleting histone H1 were distinctly different from c-MYC expression, and these changes were also inconsistent with promoting licensing^65^. Notably, we deliberately restricted our analyses to the immediate effects of c-MYC expression on licensing to minimize secondary effects from chronic c-MYC expression.

Our findings that c-MYC increases licensing speed through TRRAP, an adapter for GCN5 and multiple chromatin regulators, is consistent with prior reports of c-MYC-induced local chromatin changes^58,60^. The licensing-impaired W135E mutant deficient for TRRAP binding^26,39^ still induces histone acetylation, but does not induce histone H1 eviction at all. The minimal licensing activity of this point mutant may reflect residual TRRAP binding^26,39^ and/or alternative TRRAP-independent recruitment of histone acetylating enzymes^42^. Lysine acetyltransferases such as GCN5 and p300 also have many non-histone DNA replication substrates^66,67^, and it is possible that other substrates are particularly relevant for licensing speed control. Nonetheless, because the W135E mutation separates histone acetylation induction from histone H1 eviction, and depleting histone H1 strongly accelerated licensing, we propose that histone H1 eviction is the primary chromatin-based driver of rapid licensing.

Our findings also demonstrate that G1 length and origin licensing speed can be mechanistically uncoupled. For instance, cyclin E1 overexpression shortens G1 but fails to accelerate licensing. In addition, cells overexpressing the MB0 deletion of c-MYC have fast licensing, but no substantial change in G1 length. WT c-MYC expressing cells co-treated with siTRRAP or siCDT1 have a short G1 similar to cells with normal TRRAP or CDT1, yet licensing speed was severely diminished which promotes underlicensing. Studies by our group and others indicate that underlicensing in short G1 cells is especially detrimental for heterochromatin^8,68^. The fact that WT c-MYC-overexpressing cells are not particularly underlicensed, and c-MYC promotes MCM loading near the heterochromatin marker HP1, indicates that rapid licensing is occurring throughout the genome, including in heterochromatin. In support of this conclusion, overexpressed c-MYC has been reported to bind more than 25,000 genomic sites, many are in intergenic regions and far from promoters^69–71^, and we detect even moderately elevated c-MYC associated with heterochromatin.

Although we define a mechanism for inducing rapid origin licensing through c-MYC, we expect that alternative pathways for genome-wide chromatin opening could have similar effects. For example, there are multiple methods of reprogramming from slow-licensing somatic cells into fast-licensing pluripotent cells, some of which do not use exogenous c-MYC^72^. Interestingly, chemical reprogramming protocols have been developed from multi-step sequential signaling and chromatin modifying manipulations^73^, and we predict that during that process, origin licensing becomes rapid. G1 phase is also naturally very short in activated T cells, meaning they may also license quickly^74^, and erythroid progenitors also shorten G1 as they approach terminal differentiation^75^. Moreover, multiple oncogenes can shorten G1 when activated or over-expressed^76–78^. The very immediate effects of activating such oncogenes could include underlicensing, genome instability, and proliferation defects, whereas adapting to the perturbation presumably accelerates origin licensing^7^. Our study provides mechanistic insight into how the speed of origin licensing can be modulated through global changes in chromatin to avoid underlicensing and ensure genome stability.

## METHODS

### Cell culture

Cell lines were originally acquired from ATCC and tested for mycoplasma contamination. HEK293T cells were used for transient transfections and lentiviral packaging. RPE1-hTERT (i.e., “RPE1” or “RPE”) or HPNE cells were used for the rest of experiments. All cell lines were grown at 37°C in a 5% CO2 incubator. RPE and HPNE were cultured in DMEM (Sigma-Aldrich Chemistry) medium supplemented with 10% FBS (Sigma), 2 mM L-glutamine (Gibco), and 1x Pen-Strep (Gibco).

### Cloning

#### Plasmids encoding inducible MYC constructs

Human c-MYC bearing N-terminal double HA epitope tags was cloned into pINDUCER20 (plasmid #44012; Addgene) as previously described ^7^. Double HA tag protein sequence: YPYDVPDYAGYPYDVPDYA. The pINDUCER20-HA-HA-c-MYC plasmid was digested with *Sal*I and *BstB*I to replace the WT c-MYC open reading frame with various PCR products encoding c-MYC mutants by Gibson Assembly (New England Biolabs) following the manufacturer’s protocol. The CTD removed in *^2xHA^c-MYC^ΔCTD^*omits amino acids 355-439, while the *^2xHA^c-MYC^ΔNTD^*omits amino acids 1-143. The following three plasmids were synthesized (Twist Biosciences): ^2xHA^c-MYC ^ΔMB0^ omits amino acids 1-37 which encompasses MB0 at amino acids 18-24, ^2xHA^c-MYC with a single point mutation W135E, or ^2xHA^N-MYC were each inserted into pINDUCER20 with the cDNA of a double HA tag at the N-terminus of each of the indicated proteins. The N-MYC cDNA sequence was obtained from pMXs-hu-N-Myc (plasmid #50772; Addgene). Successful cloning was confirmed by Sanger sequencing.

#### Plasmid encoding inducible OKS (OCT4-KLF4-SOX2)

The FUW-tetO-hOKMS plasmid (plasmid#51543; Addgene) was digested with *EcoR*I to linearize it and remove the original OKMS insert. The OKSIM plasmid (plasmid#24603; Addgene) was used as a template for amplifying the OCT4, KLF4, and SOX2 open reading frames using conventional PCR. The amplified OKS fragments were assembled into the linearized FUW-tetO plasmid by Gibson Assembly (New England Biolabs) following the manufacturer’s protocol. Successful cloning was confirmed by Sanger sequencing.

#### Plasmids

- pInducer20 was a gift from Stephen Elledge (Addgene plasmid # 44012; http://n2t.net/addgene:44012; RRID:Addgene_44012) ^79^
- FUW-tetO-hOKMS was a gift from Tarjei Mikkelsen (Addgene plasmid # 51543; http://n2t.net/addgene:51543; RRID:Addgene_51543) ^18^
- OKSIM was a gift from Jose Cibelli (Addgene plasmid # 24603; http://n2t.net/addgene:24603; RRID:Addgene_24603) ^80^
- FUW-tetO-hOCT4 was a gift from Rudolf Jaenisch (Addgene plasmid # 20726; http://n2t.net/addgene:20726; RRID:Addgene_20726) ^81^
- FUW-tetO-hSOX2 was a gift from Rudolf Jaenisch (Addgene plasmid # 20724; http://n2t.net/addgene:20724; RRID:Addgene_20724) ^81^
- FUW-tetO-hKLF4 was a gift from Rudolf Jaenisch (Addgene plasmid # 20725; http://n2t.net/addgene:20725; RRID:Addgene_20725) ^81^
- Clover-Geminin(1-110) was a gift from Michael Lin (Addgene plasmid # 83915; http://n2t.net/addgene:83915; RRID:Addgene_83915) ^31^
- pFUW-tetO-L-MYC was a gift from Filipe Pereira (Addgene plasmid # 139817; http://n2t.net/addgene:139817; RRID:Addgene_139817)^82^
- pMXs-hu-N-Myc was a gift from Shinya Yamanaka (Addgene plasmid # 50772; http://n2t.net/addgene:50772; RRID:Addgene_50772)^83^

### Total lysate and chromatin fractionation

Cells were harvested via trypsinization. Fresh pellets right after harvesting or frozen pellets stored at −80°C were used for cell lysis. For total protein lysates, cells were lysed for 30 minutes on ice in standard CSK buffer (100 mM sodium chloride, 300 mM sucrose, 3 mM magnesium chloride, and 10 mM PIPES pH 7.0) with 0.5% Triton X-100 and a mixture of protease inhibitors: (1 µg/mL pepstatin A, 0.1 mM AEBSF, 1 µg/mL aprotinin and 1 µg/mL leupeptin), and phosphatase inhibitors: (1 mM β-glycerol phosphate, 10 µg/mL phosvitin and 1 mM Na-orthovanadate). Lysates were centrifuged at 4°C at maximum speed for 7.5 minutes to clarify, and supernatants were transferred to new tubes. Protein quantification using the Bradford assay (Biorad) was performed then whole cell lysates were directly diluted in SDS loading buffer to a final concentration of 1% SDS, 2.5% beta-mercaptoethanol. For SDS-PAGE gels, 20-30 μg of total protein was loaded.

Chromatin fractionation was performed as described previously ^84^. Briefly, after cell lysis for 30 minutes in CSK buffer on ice (as described above), samples were centrifuged, and the supernatant (soluble/unbound fraction) was separated from the pellet (bound fraction) by cold centrifugation for 8 minutes at 376 x g. Soluble fractions were used for protein quantification to achieve equal loading. Pellets were then cold-incubated and washed in CSK buffer two times to remove all remaining soluble proteins. Pellets were incubated for 30 minutes at room temperature in CSK buffer supplemented with 1 mM ATP, 5 mM CaCl2 and S7 nuclease (Roche) to release all the DNA-bound proteins and then centrifuged again. Protein concentrations of the supernatants were quantified then diluted in SDS loading buffer. For SDS-PAGE gels, 8 μg of chromatin-bound protein was loaded.

### Immunoblotting

Samples diluted in SDS loading buffer were boiled for 5 minutes Proteins were separated by SDS-polyacrylamide gels, followed by wet transfer onto polyvinylidene difluoride (PVDF) or nitrocellulose membranes (Thermo Fisher Scientific). Ponceau S staining of total protein (Sigma-Aldrich) was used as the loading control in all experiments. Membranes were then blocked for 1 hour at room temperature in 5% milk diluted in Tris-buffered-saline-0.1% Tween 20 (TBST). Primary antibody was diluted in 2.5% milk-TBST and incubated with membranes overnight at 4 °C. Blots were washed the next day three times with 1x TBST and incubated with the appropriate secondary antibody for 1 hour at room temperature. Blots were washed three times with 1x TBST then incubated with ECL Prime (Amersham) for 5 minutes and imaged using a Chemidoc Imaging system (BioRad).

Primary antibodies and their dilutions used for immunoblotting were the following: Cyclin E1 (1:2000, Cell Signaling Technology, 4129), CDT1 (1:1000, Cell Signaling Technology, 8064 or (1:1000, Santa Cruz Biotechnology (F-6 clone), sc-36505, OCT4 (1:3000-1:5000, Cell Signaling Technology, 2750), c-MYC (1:1000, Invitrogen (9E10 clone), MA1-980), HA (1:1000, Santa Cruz, sc-7392 clone F-7), KLF4 (1:1000-1:3000, Cell Signaling Technology, 12173), SOX2 (1:5000, Cell Signaling Technology, 3579), CDC6 (1:1000, Cell Signaling Technology, 3387), E2F1 (1:1000, Cell Signaling Technology, 3742), Cyclin A2 (1:2000, Cell Signaling Technology, 4656), L-MYC (1:1000, Cell Signaling Technology, 76266), H1.0 (1:1000, Abcam, ab11079), HP1 (1:2000, Santa Cruz Biotechnology, sc-515341), H3K9me3 (1:2000, Activ motif, 39062), pan-H1 (1:2000, Santa Cruz Technology, sc-8030 clone AE-4), MCM2 (1:10,000, BD Biosciences, 610700), MCM6 (1:1000, Santa Cruz Biotechnology, sc-9843), ORC2 (1:2000, Cell Signaling Technology, 4736, 3G6), ORC4 (1:3000, Santa Cruz Biotechnology, 136331). Secondary antibodies used were: donkey anti-rabbit IgG HRP-conjugated (1:10,000, Jackson ImmunoResearch, 711-035-152), donkey anti-mouse IgG HRP-conjugated (1:10,000, Jackson ImmunoResearch, 715-035-150), anti-rat IgG HRP-conjugated (1:10,000, Jackson Immunoresearch).

### Quantitative flow cytometry

To label cells that are actively synthesizing DNA, cultures were incubated with 10 μM EdU (Santa Cruz Biotechnology) for 30 minutes prior to harvesting. Chromatin-binding was carried out essentially as described in ^6^. Briefly, cells were harvested and soluble proteins removed by extraction on ice for 8 minutes using CSK buffer: (100 mM sodium chloride, 300 mM sucrose, 3 mM magnesium chloride, and 10 mM PIPES pH 7.0) supplemented with 0.5% triton X-100 and a mixture of protease and phosphatase inhibitors as above. Cells were washed with 1% BSA in phosphate-buffered saline (PBS) and fixed in 4% paraformaldehyde (Sigma-Aldrich Chemistry) diluted in PBS for 15 minutes at room temperature. Cells were washed in 1% BSA-PBS and stored at 4 °C until staining. Total protein flow cytometry was performed without pre-extraction with CSK. Briefly, after cells were trypsinized and washed with 1x PBS, cells were directly fixed with 4% paraformaldehyde in PBS for 15 minutes at room temperature. Then, samples were washed with 1% BSA-PBS and stored overnight at 4°C. Fixed cells were then permeabilized with 0.5% Triton x-100 in 1% BSA-PBS at room temperature for 15 min, then 1% BSA-PBS was added to pellet cells before EdU labelling.

For EdU detection, cells were incubated in EdU labeling solution: (1 μM Alexa-fluor 647 or 488 azide (Life Technology), 1 mM CuSO4, and 100 mM ascorbic acid diluted in PBS) for 30 minutes at room temperature in the dark. Cells were washed in 1% BSA-PBS + 0.1% NP-40 solution. For primary antibody staining, cells were incubated with the designated primary antibody diluted in 1% BSA-PBS + 0.1% NP-40 for 1 hour at 37°C in the dark then washed with 1% BSA-PBS + 0.1% NP-40 solution. For secondary antibody staining, cells were incubated with the designated secondary antibody for 1 hour at 37°C in the dark then washed with 1% BSA-PBS + 0.1% NP-40 solution. Finally, cells were incubated in 1 μg/mL DAPI (Sigma-Aldrich Chemistry) and 100 μg/mL RNase (Sigma-Aldrich Chemistry) diluted in 1% BSA-PBS + 0.1% NP-40 for 1 hour at 37°C in the dark or overnight at 4 °C. Data were collected the next day using an Attune NxT Flow cytometer. Control samples were not incubated in EdU labeling solution or primary antibody to determine thresholds for positive signal.

Primary antibodies and their dilutions used for flow cytometry were the following: MCM2 (1:190, BD biosciences, 610700), ORC4 (1:50, Santa Cruz Biotechnology, 136331), CDT1 (1:100, Abcam, (EPR17891) ab202067), H3K14Ac (1:200, Millipore, 07-353, LOT: 2302183), H3K9Ac (1:200, Millipore, 07-352), H1X (1:100, ThermoFisher, A304-604A), ɣH2AX (1:200, Cell Signaling Technology, 9718S), ORC6 (1:50, Santa Cruz Biotechnology, sc-32735). Secondary antibodies used were: donkey anti-mouse secondary antibody conjugated to Alexa-fluor 488 or Alexa-fluor 594 for MCM2 or Alexa-fluor 488 for ORC4, or donkey anti-rabbit secondary antibody conjugated to Alexa-fluor 647 for CDT1, H3K9Ac, H3K14Ac, H1X, ɣH2AX, or anti-rat secondary antibody conjugated to Alexa-fluor 647 for ORC6 (1:1000, Life Technology).

### Statistics

The number of independent biological replicates from cells plated and analyzed on different days are shown by symbols on the box plots (replicate 1 = closed circle, replicate 2 = closed triangle, replicate 3 = open circle, replicate 4 = open triangle). To avoid pseudoreplication ^85^, statistical testing was only performed using the median values, and not individual cell measurements.

#### Flow cytometry data analysis

Data analysis was performed as described in ^6^. FCS data files were analyzed in FCS Express 7 software (De Novo). Each sample within an experiment was randomly downsampled to the same number of cells. Raw single cell data values were exported from FCS Express to GraphPad Prism. Statistical analyses were performed using GraphPad Prism version 10.6.1. For flow cytometry experiments, thousands of single-cell events were collected per sample, and the Median Fluorescence Intensity (MFI) was calculated for each condition.

Because flow cytometry data are log-normally distributed and subject to day-to-day experimental variance (batch effects), Ratio Paired *t*-tests were executed on the raw MFI values. This approach automatically log-transforms the data to accurately assess the multiplicative shift between the control and treatment groups while preserving individual replicate variance. Outliers were removed using ROUT (Q = 2%), and *p*-values are reported in figure captions.

Raw signal intensities between replicate experiments, but not between samples within an experiment, were affected by occasional voltage changes for the flow cytometer. In these cases, median intensities were normalized to each respective control and shown in figures next to a representative experiment.

### Fluorescence Activated Cell Sorting (FACS)

#### Pre-sorting validation

To validate the expression of the ELDR-Glo (mCherry variant) ^30^ and Gem1-110 (mClover) ^31^ biosensors by flow cytometry as in Supplementary Fig. S8, cells were harvested by trypsinization, washed once with 1x PBS, then resuspended in FluoroBrite^TM^ DMEM (Invitrogen) supplemented with 5% FBS, 2 mM L-glutamine and 1x Pen-Strep. The Vybrant Dye Cycle Violet stain (Invitrogen, cat# V35003) was added as the DNA dye to the cell suspension at 1:1000, mixed well and incubated for ∼1 hour at 37°C. Cells were kept at 37°C until right before sorting the samples on the Attune NxT Flow analyzer. The ELDR-Glo biosensor was detected by the mCherry channel, and the Gem1-110 biosensor by the Alexa-fluor 488 channel.

#### Pre-sorting experimental setup

Cells were plated three days before the sorting in a 15-cm cell culture plate. The number of plated cells was adjusted according to the final desired number of sorted cells. Control cells (parental RPE1 cells with or without DNA dye, or cells with only ELDR-Glo and DNA dye) were plated in 6-cm plates for gating purposes only. 20 ng/mL of doxycycline was added 24 hours before sorting to the + c-MYC condition. On the day of sorting, cells were harvested by trypsinization, filtered through 50 μm Celltrics filters (Sysmex) to discard clumps and debris. Cells were counted, washed once with warm 1x PBS, then resuspended in FluoroBrite^TM^ DMEM (Invitrogen) supplemented with 10% FBS, 2 mM L-glutamine and 1x Pen-Strep. The Vybrant Dye Cycle Violet stain (Invitrogen, cat# V35003) was added as the DNA dye to the cell suspension, mixed well and incubated for ∼1 hour at 37°C. Cells were kept at 37°C until right before sorting.

#### Cell Sorting and pellet processing

Cell sorting was performed as five-way sorting on the FACSymphony S6 at room temperature. Gating was determined according to the control cells as described above. Cells were separated into early G1, mid G1, late G1, G0, S/G2/M populations. After sorting was complete, sorted and unsorted cells were held on ice, pelleted, washed with 1x PBS, then flash frozen and stored at −80°C for later use.

### siRNA transfection

si-Luciferase (si-LUC - control siRNA) or treatment siRNA were transfected into cells using the Lipofectamine RNAiMAX (Invitrogen) reagent according to the manufacturer’s instructions. Briefly, target RPE1 cells were plated on Day 0. On Day 1, siRNA or Lipofectamine were first mixed in Opti-MEM (Gibco) medium individually. Then, both mixtures were added together and incubated for 15 minutes. The original culture medium was aspirated and replaced with fresh antibiotic-free DMEM with L-glutamine at a final 10% FBS concentration. Then the mixture containing the siRNA and Lipofectamine was added to cells. Doxycycline was added 12 hours (for the si-CDT1 sample) or 24 hours (for the si-TRRAP or si-H1X samples) later, and samples were collected 24 hours later for a total of 36 hour for si-CDT1 or 48 hour siRNA treatment for si-TRRAP or si-H1X. si-LUC: 20 or 100 nM, 5′-UCGAAGUACUCAGCGUAAG-3′; si-TRRAP: 20 nM, 5’-AAUGGCUCGAAAGGAAUGG-3’; si-H1X: 100 nM, 5′-CCAAGAAGGUUCCGUGGUUTT-3′; si-CDT1: 100 nM, 5′-CCUACGUCAA GCUGGACAATT-3′

### Cell line generation

To generate cells stably expressing proteins of interest, lentivirus stocks were produced by co-transfecting HEK293T cells with lentiviral expression plasmids in addition to ΔNRF and VSVG virus packaging plasmids (gifts from J. Bear) using 50 μg/mL polyethylenimine (PEI)-Max (Sigma-Aldrich Chemistry). Virus-containing supernatants were collected, filtered and in some cases concentrated using Lenti-X Concentrator (Clontech, cat. #631231). Collected viral supernatants were used to transduce target cells: RPE1, HPNE, or RPE1 cells constitutively expressing rtTA (reverse tetracycline Trans Activator) using 8 μg/mL polybrene (Millipore) for 24 hours. In some cases, a second round of viral transduction was performed to ensure sufficient expression. Drug selection was performed with 500 μg/mL geneticin (Gibco) for 5 days, when appropriate. To generate monoclonal cell lines, cells were plated following drug selection for clonal selection, and clones were screened for positive expression of the construct of interest by immunoblotting or IF imaging.

### Chemical reagents/inhibitors

These inhibitors/chemicals were used in this study: **Doxycycline (Dox)** (CalBiochem, cat. #32485) at the indicated concentrations in each figure caption, **Trichostatin A (TSA)** (histone deacetylase inhibitor) (Sigma Aldrich) at 300 nM or 1 μM, **Butyrolactone (MB-3)** (GCN5 inhibitor) (Santa Cruz Biotechnology, cat. #sc-358657) at 200 μM, **NU9056** (TIP60 inhibitor) (Abcam, cat. #ab255734) at 25 μM, **WM-3835** (HBO1 inhibitor) (Sigma, cat. #SML3319) at 5 μM, **C646** (p300 inhibitor) (Sigma, cat. #382113) at 10 μM, **Hydroxyurea** (HU) (Selleck, cat. #S1896) at 2 mM.

### Proximity ligation assay (PLA)

The PLA assays were performed according to the Duolink PLA kit manufacturer’s protocol (Sigma) with minor modifications, as described previously ^86^. Briefly, cells were pulsed with EdU, then pre-extracted using CSK + 0.5% Triton-X, prior to fixing in 4% methanol-free formaldehyde for 10 minutes at RT. EdU labeling was performed as described previously for flow cytometry. Cells were blocked in a solution of 5% normal goat serum, 5% BSA and 5% sucrose in PBS for 30 minutes, followed by primary antibody incubation in blocking solution for either 1 hour at RT or overnight at 4°C. The cells were washed 3x for 10 minutes each using wash buffer A (10 mM Tris pH 7.5, 150 mM NaCl, 0.05% Tween-20) and treated with plus and minus probes diluted in blocking solution for 1 hour at 37°C. After two washes with buffer A, the probes were ligated for 30 minutes at 37°C, followed by *in situ* PCR amplification using fluorescent oligonucleotides (green) for 120 minutes at 37°C. Cells were washed 2x for 10 minutes each in buffer B (200 mM Tris pH 7.5, 100 mM NaCl). DNA was counterstained with DAPI and mounted using ProLong Gold Antifade mounting medium (Life Technologies). Images were acquired on a Zeiss 800 confocal microscope with 40X oil-immersion objective. Images were processed using Fiji/ImageJ, and PLA pairs were quantified using CellProfiler (version 4.2.4). Primary antibodies used for PLA are: c-MYC (1:250, Abcam, clone# Y69), HP1 (1:500, Santa Cruz Biotechnology, sc-515341), and MCM3 (1:500, Bethyl laboratories, A300-192A).

### PLA analysis

PLA foci were quantified using a previously published pipeline from cellprofiler.org, with minor modifications. Briefly, nuclei were segmented based on DAPI signal. S-phase cells were identified based on EdU signal and masked onto the DAPI objects to eliminate S-phase cells from downstream analyses. G2 cells were characterized by lack of PLA signal due to low levels of chromatin-bound MCM in G2 cells ^14^. PLA objects were detected with a 4-10 pixel limit and global thresholding. PLA objects were related to the masked nuclei, and per-object counts of PLA foci were calculated and exported. PLA spot counts were analyzed on a per-nucleus basis. For each condition, empirical cumulative distribution functions (ECDF(x) = % nuclei with PLA foci ≤ x) were generated by calculating the percentage of nuclei with PLA spot counts less than or equal to each threshold. ECDFs were calculated separately for each biological replicate, then plotted as the mean ± SEM across replicates.

### Immunofluorescence (IF)

For IF imaging of the ELDR-Glo and the Gem1-110 biosensors, RPE1 cells expressing both biosensors were cultured in multi-well glass-bottomed dishes. Cells were fixed with 4% paraformaldehyde for 15 minutes at room temperature, then washed with 1x PBS. Cells were imaged with a 20x objective lens (Plan-Apo 20x/0.8 WD 0.55mm) on a Zeiss 880 confocal microscope, and the images were acquired with the ZEN acquisition software (Zeiss). No photobleaching was observed during image acquisition. Images were analyzed using Fiji (ImageJ).

For IF imaging of H3Ac, HP1, and H1X, RPE1 cells were pulsed with 10 µM EdU for 20 minutes prior to pre-extraction in CSK+0.5% Triton-X, followed by fixation for 10 minutes at RT using 4% methanol-free formaldehyde in PBS. Cells were incubated in blocking solution containing 5% BSA and 2% normal goat serum in PBS for 30-45 minutes at RT. Cells were incubated with primary antibodies diluted in blocking solution for either 1 hour at RT or overnight at 4°C. Cells were subsequently washed with PBS-0.05% Tween-20 three times for 10 minutes each, followed by secondary antibody incubation for 1 hour at RT. DNA was counterstained with DAPI. Representative images were acquired on either Zeiss LSM 880 or LSM 800 using a 63X oil immersion objective. All images were brightness/contrast-adjusted equally for clarity.

Primary antibodies used for IF were: HP1 (1:500, Santa Cruz Biotechnology, sc-515341), *pan-*H3Ac (1:1000, Millipore Sigma, 06-599), and H1X (1:250, ThermoFisher, A304-604A). Secondary antibodies used were: donkey anti-mouse secondary antibody conjugated to Alexa-fluor 488 for HP1, or donkey anti-rabbit secondary antibody conjugated to Alexa-fluor 488 for pan-H3Ac or Alexa-fluor 594 for H1X (1:1000, Life Technology).

### H3Ac imaging and analysis

Images were acquired on GE INCELL 2200 widefield high-content imager using a 20X air objective under identical microscope settings for all conditions within each experiment. Images were acquired from multiple fields of view from independent biological replicates. Images were then analyzed using CellProfiler. Nuclei were segmented based on DAPI signal. S-phase cells were identified based on EdU signal and masked onto the DAPI objects to eliminate S-phase cells from downstream analyses. H3Ac signal was related to the masked DAPI objects. Integrated mean intensities per nucleus of H3Ac were quantified and exported.

### Live-cell imaging

RPE1 cells were plated on glass-bottom plates (Cellvis) in regular DMEM and media was switched to FluoroBrite^TM^ DMEM (Invitrogen) supplemented with 10% FBS, 2 mM L-glutamine and 1x Pen/-Strep right before imaging. Nikon Ti Eclipse inverted microscope was used for fluorescence time-lapse imaging of a total of 48-72 hour movie using Plan Apochromat dry objective lenses 20x (numerical aperture (NA) 0.75). Images were captured using an Andor Zyla 4.2 sCMOS detector with a 12-bit resolution every 20 min. Autofocus was performed by the Nikon Perfect Focus System. To maintain temperature at 37°C and inside atmosphere at 5% CO2, a custom enclosure humidified chamber (Okolabs) was used. The filter set used was from Chroma: CFP - 436/20, 455 (excitation, beam splitter and emission filter). Every 10 minutes, images were collected using the NIS-Elements AR software. No phototoxicity was generally observed however, with high c-MYC expression at longer time points, cell death was observed in imaging locations but not in other non-exposed locations or in control cells.

### Segmentation, tracking and PCNA analysis

Cell segmentation and mask generation were performed using Cellpose ^87^. We extracted cell features using “regionprops” from scikit-image. Manual corrections to segmentation and object linking were done in napari through a custom-built interface. For cell tracking, we used btrack ^88^. Cells were tracked right at the start of mitosis. Once PCNA foci started to appear or disappear, S phase start or end was noted, respectively. Blurry cells or cells not showing a clear PCNA signal for transitions between phases were manually removed from further analysis.

### qPCR

RNeasy plus mini kit (Qiagen, cat # 74134) was used to extract total RNA from cells. High-Capacity Reverse Transcription kit (Thermo Fisher Scientific, cat # 4368814) was used to reverse transcribe the RNA to cDNA according to manufacturer’s instructions. iTaq Universal SYBR Green Supermix (Biorad, cat # 1725121) was used for qPCR on the QuantStudio 7 system (Thermo Fisher Scientific) using the standard sybergreen protocol. Each sample was pipetted in technical duplicates for each analyzed gene. To calculate mRNA abundances: the 2^^-ΔΔCt^ method was used to normalize the average Ct value of a given gene to the average Ct value of ACTB (housekeeping gene), then normalize to the untreated cells as the experimental biological control sample.

The following primer sequences were used in this study: ***CDT1***, 5’-GTTTCCAGCCCCAGTACC-3’ (fwd) and 5’-ATGACGCAAGCTCAGAGATG-3’ (rev); ***CDT1-2***, 5’-GTAGGCGTTTTGAGGAGTGC-3’ (fwd) and 5’-TAATCTGACCTCCTGGTGCC-3’ (rev); ***MCM2***, 5’-ATTTCGTCCTGGGTCCTTTC-3’ (fwd) and 5’-CGCTGGTAGTTCTGATAGATGG-3’ (rev); ***CCNA2***, 5’-CGCTGGCGGTACTGAAGTC-3’ (fwd) and 5’-GAGGAACGGTGACATGCTCAT-3’ (rev); ***E2F1***, 5’-TCTCCGAGGACACTGACAG-3’(fwd) and 5’-ATCACCATAACCATCTGCTCTG-3’ (rev); ***CCNE1***, 5’-TGCAGCCAAACTTGAGGAAATC-3’ (fwd) and 5’-TAGTCAGGGGACTTAAACGCCA-3’ (rev); **MCM*6***, 5’-GTGATCAGGGATGTAGAACAGC-3’ (fwd) and 5’-AGCTTGGGTCTCTTGAATACG-3’ (rev); ***ACTB***, 5’-ACCTTCTACAATGAGCTGCG-3’ (fwd) and 5’-CCTGGATAGCAACGTACATGG-3’(rev).

### RNA sequencing

#### Sample preparation

RPE1 cells were treated with 20 ng/mL doxycycline for 24 hours prior to harvesting to express ectopic WT or W135E c-MYC. Cells were harvested, counted, washed with 1x PBS, then resuspended in Zymo 1X DNA/RNA shield (Zymo Research, cat # R1100-50) according to Plasmidsaurus instructions. Samples were shipped at room temperature. Total number of samples was sixteen, as follows: RPEs -/+ WT c-MYC (4 replicates each) and RPEs -/+ W135E c-MYC (4 replicates each).

#### Sequencing and preprocessing

RNA-seq (3’ Tag-seq) of polyA+ transcripts was performed by Plasmidsaurus on an Illumina NovaSeq X Plus to achieve ∼ 10 million deduplicated, single-end ∼90-bp reads per sample. Raw sequencing reads were processed by Plasmidsaurus. Briefly, reads were quality-filtered using fastp v0.24.0 with poly-X tail trimming, 3’ quality-based tail trimming, a minimum Phred quality score of 15, and a minimum length requirement of 50 bp. Quality-filtered reads were aligned to the Ensembl GRCh38 (release 114) reference genome using STAR v2.7.11 ^89^ with non-canonical splice junction removal. PCR and optical duplicates were removed using UMI-based deduplication with UMIcollapse v1.1.0. Gene-level quantification was performed using subread featureCounts v2.1.1^90^ with strand-specific counting, multi-mapping read fractional assignment, exons and 3’ UTR as the feature identifiers, and grouped by gene ID.

#### Differential Expression Analysis

We performed differential gene expression (DGE) analysis using DESeq2^91^ in R v4.5.3, filtering for protein-coding genes with ≥10 counts in at least 4 samples. Fractional counts were rounded to the nearest integer. The generalized linear model adjusted for genotype (type: full-length [WT] vs. mutant [W135E] MYC), treatment (condition: control vs. doxycycline), and their interaction, using WT and control as reference levels. Log2 fold changes (LFC) were shrunk using the ashr^92^ estimator and p-values were adjusted for multiple testing using the Benjamini-Hochberg^93^ (BH) procedure to control the False Discovery Rate (FDR).

#### Comparison to public data

To validate our findings, raw RNA-seq read counts for MCF10A cells were obtained from GEO (GSE197330)^26^. Differentially expressed genes were identified using DESeq2, filtering for protein-coding genes with ≥10 counts in at least 3 samples. Because WT and mutant MYC shared a common control group, the generalized linear model only adjusted for genotype. Direct contrasts were extracted to evaluate both genotypes (WT and W135E) against the control, as well as the W135E mutant directly against WT MYC.

#### Functional enrichment

Gene set enrichment analysis was performed using clusterProfiler ^94^ on genes ranked by LFC. Hallmark gene sets were obtained from MSigDB ^95^ using msigdbr and Biological Process and Molecular Function gene sets were obtained using org.Hs.eg.db. Gene sets were restricted to those containing between 10 and 500 genes. Significance was defined by a BH FDR-adjusted p-value < 0.05.

## Supporting information

Fleifel Supplemental Figures S1-S18

## DATA AVAILABILITY

All data are available in the main text or the supplementary materials. The RNA-seq profiling data are also available as GEO accession GSE349348.

## ACKNOWLEDGMENTS

The authors are grateful to the members of the Cook lab and many colleagues for advice and helpful discussions and to Jeffrey Jones for managerial assistance. We thank Priya Hibshman, and the Der lab, for providing the HPNE cells. We thank Juanita Limas for providing the RPE1 cell line expressing c-MYC. We thank the staff of the UNC Flow Cytometry Core Facility for assistance with cell sorting. RNA-sequencing data from Hinds, *et al*., ^26^ was accessed from the Gene Expression Omnibus under accession number GSE197323. Cartoons and illustrations were created in BioRender.

## Funding statement

NIH National Institute of General Medical Sciences (NIGMS) grant R35GM141833 (JGC).

NIH National Cancer Institute (NCI) grant R61CA302625 (JGC).

American Heart Association (AHA) 23PRE1027147 (DF).

UNC Graduate School Dissertation Completion Fellowship (A.A.A).

UNC SOM Core Facility Advocacy Committee and Office of Research Technologies (DF). The UNC Flow Cytometry Core Facility (RRID:SCR_019170) is supported in part by NIH P30 CA016086 Cancer Center Core Support Grant to the UNC Lineberger Comprehensive Cancer Center.

Research reported in this publication was supported in part by the North Carolina Biotech Center Institutional Support Grant 2017-IDG and by the NIH 1UM2AI30836-01.

## AUTHOR CONTRIBUTIONS

Conceptualization: D.F. and J.G.C. developed the research plan and conceptualization of the results through ongoing discussions. A.K. and A.A.A contributed to result conceptualization.

Methodology, investigation and visualization: D.F. performed all the flow cytometry, molecular biology experiments, and cell line generation. A.K. performed the PLA and IF imaging experiments. A.A.A. performed the RNAseq analysis and its data visualization. A.A., S.Q. and E.E. contributed to the immunoblotting experiments and live-cell imaging analysis.

Funding acquisition: J.G.C. applied for lab-level funding to support this project. D.F. applied to and was awarded a pre-doctoral fellowship and a UNC core facility voucher award.

Writing: D.F. and J.G.C. wrote and edited the original manuscript. A.K. and A.A.A contributed to the reviewing of the manuscript.

## COMPETING INTERESTS DECLARATION

Authors declare that they have no competing interests.

## ADDITIONAL INFORMATION

### Supplementary Information is available for this paper

Supplementary Figures. S1 to S18

Supplementary Tables S1 to S11

## Notes

### Competing Interest Statement

The authors have declared no competing interest.

