## Supplementary material for "Molecular control of replication licensing speed by c-MYC": Fleifel Supplemental Figures S1-S18

**SUPPLEMENTARY FIGURES S1-S18**

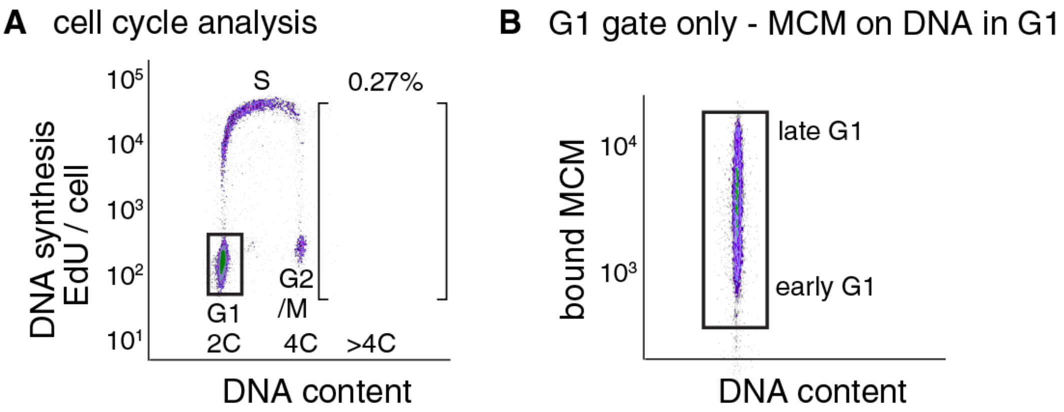


**Supplementary Fig. S1.**

**Gating scheme for analysis of cell cycle distribution and MCM loading in G1 cells.**

**(A)** Cell cycle analysis for control RPE1 cells. Cells were stained for EdU after pulse-labeling to identify S phase cells, and DAPI for DNA content. G1 cells are EdU-negative and have 2C DNA content, S phase cells are EdU-positive and have between 2C and 4C DNA content, and G2/M cells are EdU-negative and have 4C DNA content. Cells with >4C DNA content are typically the result of re-replication.

**(B)** Chromatin flow cytometry. Cells were stained for bound MCM2 with anti-MCM2 and with DAPI for DNA content. Bound MCM antibody signal intensity is shown for cells in only the G1 gate from panel (A).

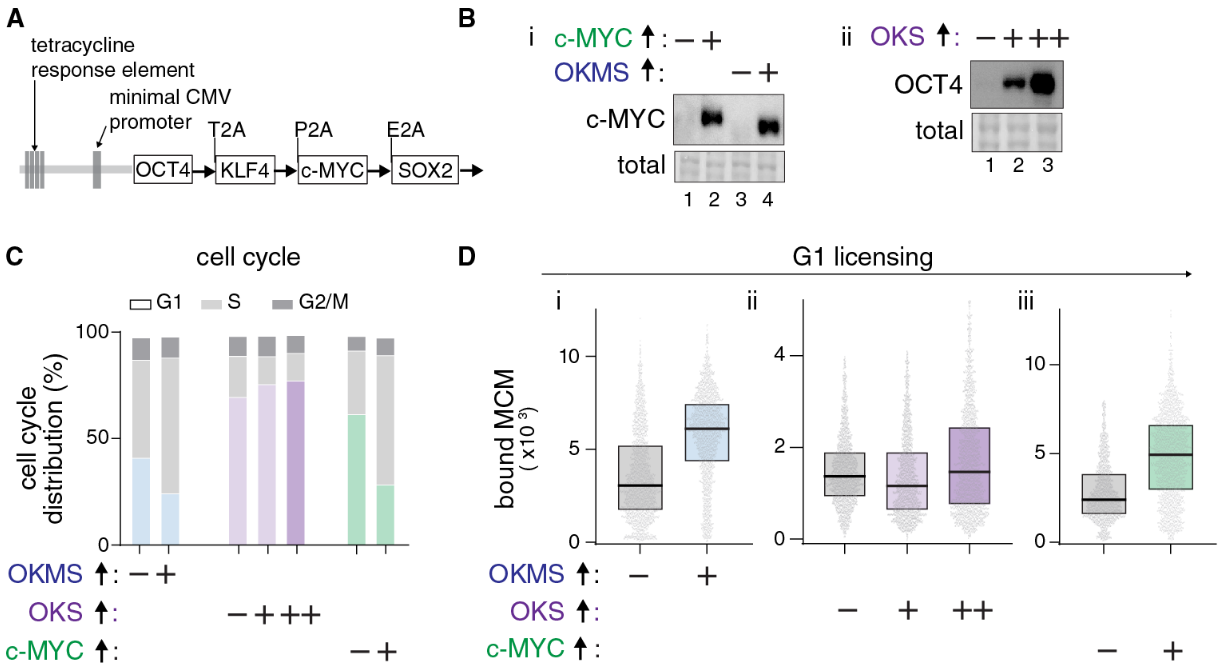


**Supplementary Fig. S2.**

**c-MYC or OKMS expression, but not OKS, accelerates licensing.**

**(A)** Schematic of the polycistronic construct from Cacchiarelli *et al.*[18].

**(B)** RPE1 cells were transduced with the construct in (A) and an rtTA-expressing construct to express the reverse tetracycline-controlled transactivator (rtTA). Immunoblotting for ectopic c-MYC and $^{2xHA}$c-MYC (i) or OCT4 (ii) as a representative of the OKS proteins in control or treated RPE1 cells overexpressing OKMS (20 ng/mL doxycycline "+") or c-MYC only (20 ng/mL"+") or OKS (20 "+", or 100 "++" ng/mL) for 24 hours, respectively; "-" indicates no doxycycline treatment. OKMS: OCT4, KLF4, c-MYC, and SOX2. OKS: OCT4, KLF4, and SOX2.

**(C)** Cell cycle phase distribution from the same samples in (B). Representative of n>3 for c-MYC or OKMS, n=1 for OKS.

**(D)** Chromatin-bound MCM per cell in G1 from the same samples in (B). 3000 cells per sample. Fold-change in median MCM intensity relative to each respective control: OKMS (i) = 2.00; OKS (ii) = 0.87 (+) and 1.09 (++); c-MYC (iii) = 1.97. Representative of n>3 for c-MYC or OKMS, n=1 for OKS.

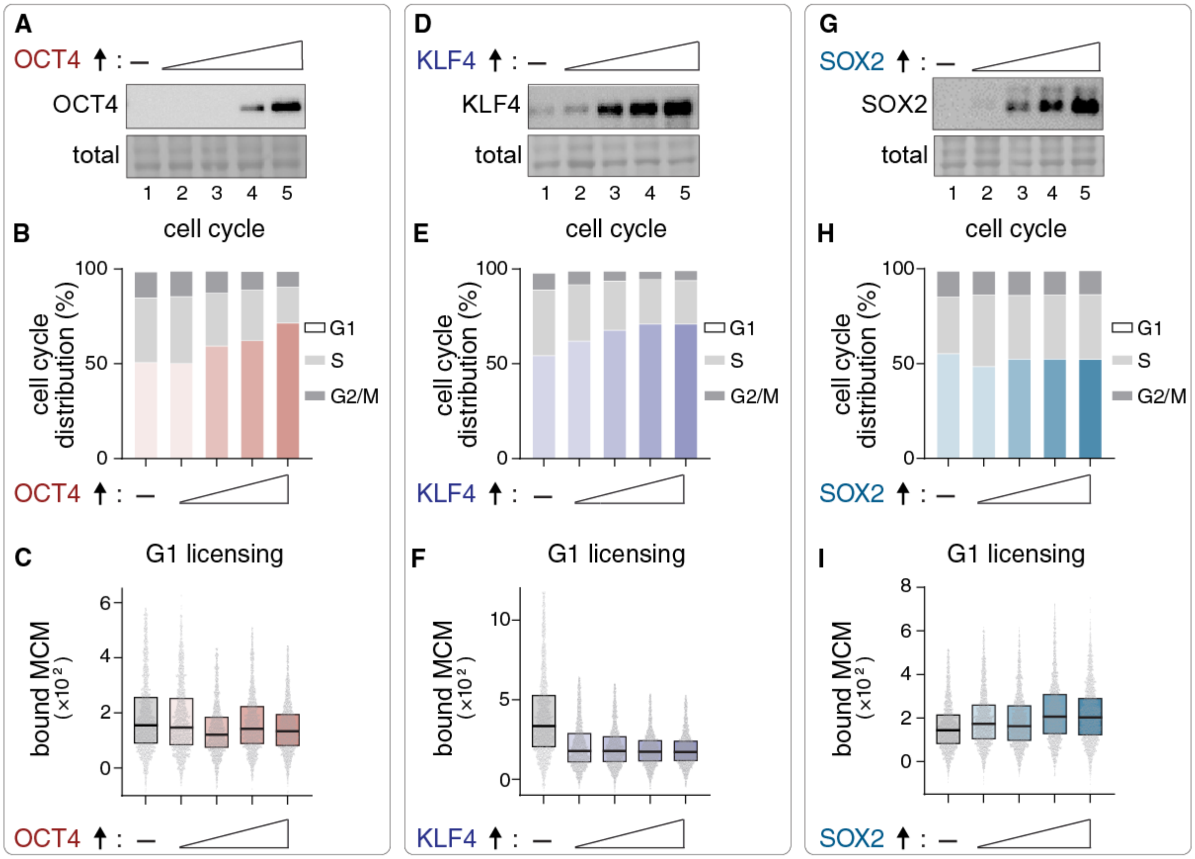


**Supplementary Fig. S3. Expressing SOX2, OCT4, or KLF4 does not accelerate licensing or shorten G1.**

**(A)** Immunoblotting for ectopic OCT4 in control or treated RPE1 cells overexpressing OCT4 at increasing concentrations of doxycycline (2, 5, 7, 10 ng/mL) for 48 hours.

**(B)** Cell cycle phase distribution of the same conditions as in (A).

**(C)** Chromatin-bound MCM per cell in G1 from the same samples in (B). 2000 cells per sample. Representative of n=1.

**(D)** Immunoblotting for ectopic KLF4 after induction with doxycycline (2, 5, 7, 10 ng/mL) for 48 hours.

**(E)** Cell cycle phase distribution of the same conditions as in (D).

**(F)** Chromatin-bound MCM per cell in G1 from the same samples in (E). 2000 cells per sample. Representative of n>3.

**(G)** Immunoblotting for ectopic SOX2 after induction with doxycycline (2, 5, 7, 10 ng/mL) for 48 hours.

**(H)** Cell cycle phase distribution of the same conditions as in (G).

**(I)** Chromatin-bound MCM per cell in G1 from the same samples in (H). 2000 cells per sample. Representative of n=1.

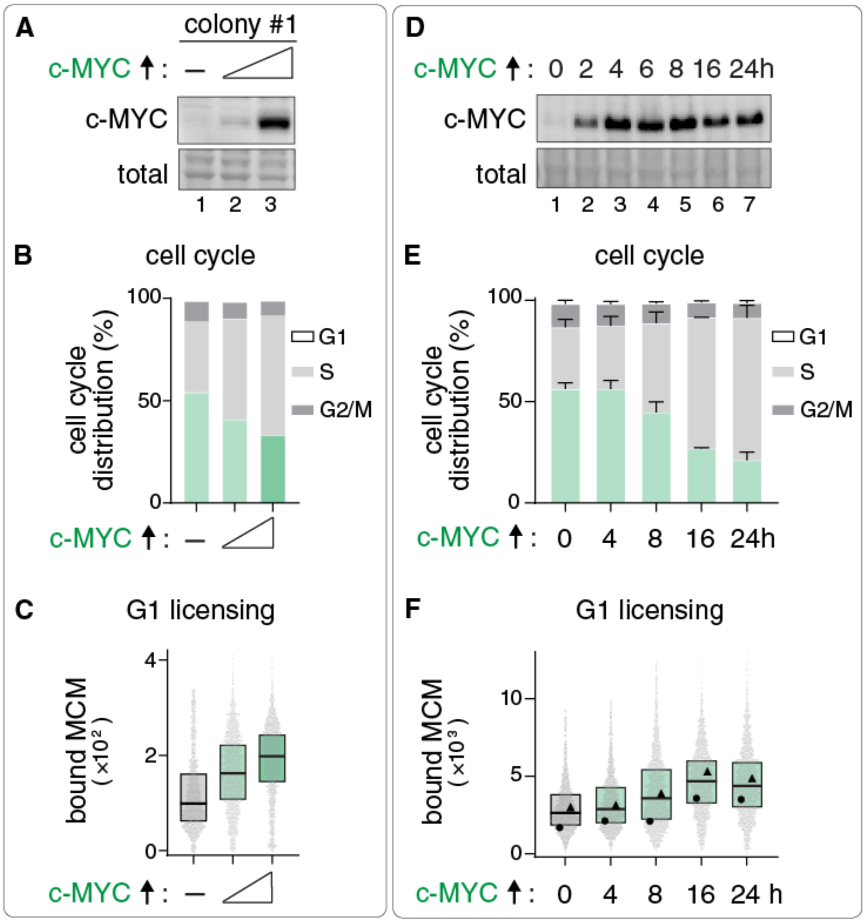


**Supplementary Fig. S4**

**c-MYC expression accelerates licensing and shortens G1 in a dose- and time-dependent manner.**

**(A)** Immunoblotting for ectopic $^{2xHA}$c-MYC in control or doxycycline-treated RPE1 cells overexpressing c-MYC at (20 and 100 ng/mL) for 24 hours. Cell line is from an independent colony separate from the cell line in Fig. 1 (colony #6).

**(B)** Cell cycle phase distribution of the same conditions as in (A).

**(C)** Chromatin-bound MCM per cell in G1 from the same samples in (B). 2000 cells per sample (n=1).

**(D)** Immunoblotting for ectopic $^{2xHA}$c-MYC in control or doxycycline-treated RPE1 cells overexpressing c-MYC at the indicated timepoints at 20 ng/mL doxycycline.

**(E)** Cell cycle phase distribution of the same conditions as in (D). Data are combined from two biological replicates; mean with error bars ± SEM.

**(F)** Chromatin-bound MCM per cell in G1 from the same samples in (E). The aggregate of two independent biological replicates was randomly downsampled to 3000 cells per sample.

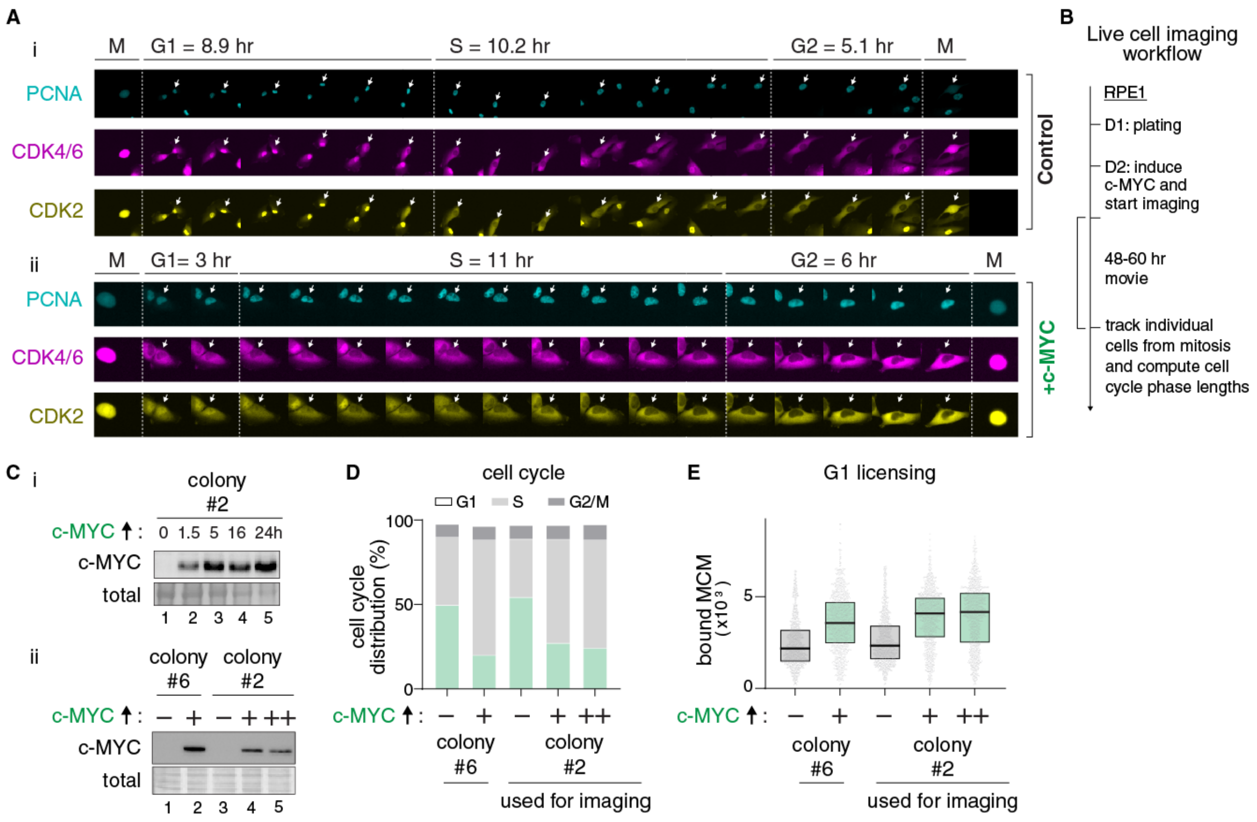


**Supplementary Fig. S5**

**Live-cell imaging, workflow, and additional validation of fast licensing.**

**(A)** Film strip of representative control (i) or c-MYC-expressing (ii) cells from wide-field time-lapse imaging (colony #2). Images were collected every 10 minutes on the microscope. Selected frames are shown every 1.27 hour or 1 hour from control or treated conditions, respectively to show an equal number of frames for both cells.

**(B)** Workflow for time-lapse live-cell imaging to compute G1 length. RPE1 cells were treated with 100 ng/mL of doxycycline (colony 2 immunoblot in panel C) right before imaging to overexpress c-MYC or left untreated (control cells).

**(C) (i)** Immunoblotting for ectopic $^{2xHA}$c-MYC in control or doxycycline-treated RPE1 cells overexpressing c-MYC using 100 ng/mL at the indicated timepoints. **(ii)** As in (i) but using 20 ng/mL (colony #6), or 50, 100 ng/mL "+", "++" (colony #2) for 24 hours, respectively.

**(D)** Cell cycle phase distribution from the same samples in (C-ii).

**(E)** Chromatin-bound MCM per cell in G1 from the same samples in (C-ii). 1000 cells per sample. Representative of n=1.

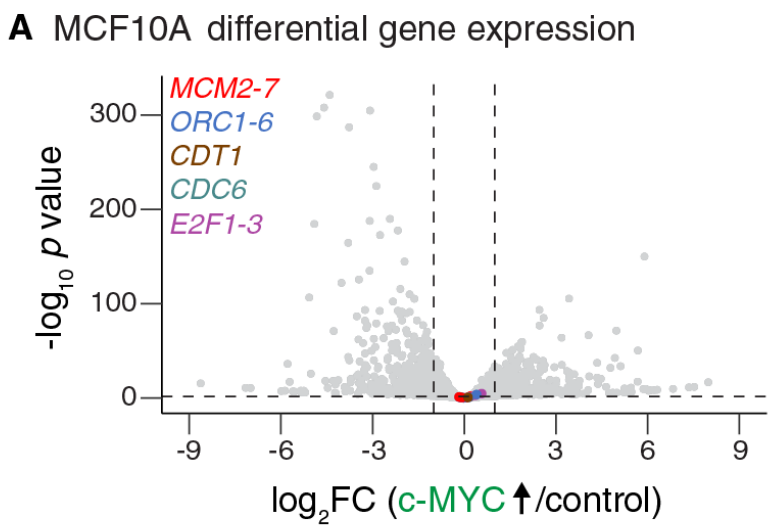


**Supplementary Fig. S6**

**c-MYC overexpression does not significantly induce licensing or E2F activator genes in MCF10A cells.**

Publicly-available data analyzed from Hinds *et al.* [26]. Volcano plot showing the differentially expressed genes in MCF10A cells constitutively-expressing WT c-MYC versus empty vector control. Origin licensing factors and E2F activator genes are color-coded. $Log_2$(fold change) threshold = ±1, adjusted *p*-value threshold = 0.05.

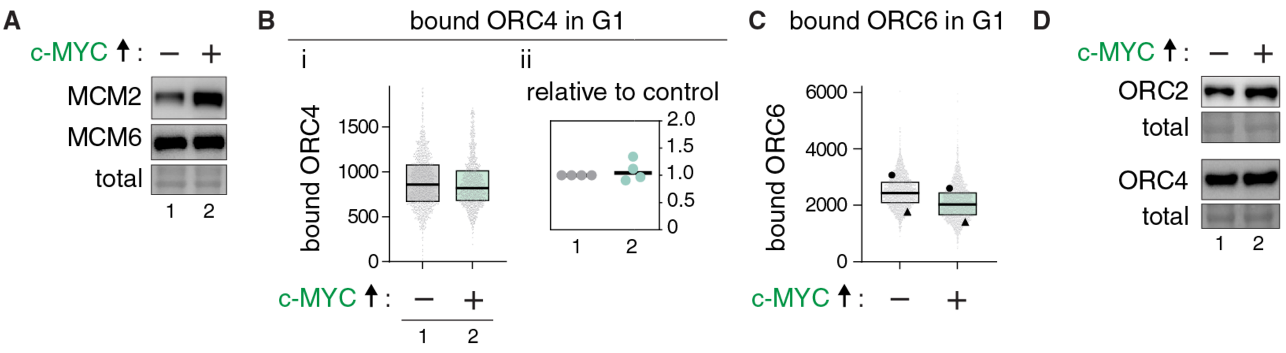


**Supplementary Fig. S7. Total MCM and ORC subunit levels or chromatin-bound ORC in G1 cells.**

**(A)** Immunoblotting for MCM2 and MCM6 in control or doxycycline-treated RPE1 cells at 20 ng/mL for 24 hours. Representative of n=2 for MCM2, n=1 for MCM6.

**(B) i-** Chromatin-bound ORC4 per cell in G1 from control or treated RPE1 cells overexpressing c-MYC at 20 ng/mL for 24 hours, representative of n=3, 1000 cells / sample. Statistical significance was determined by a ratio paired *t*-test on the median fluorescence intensities from four independent biological replicates; p=0.48. **ii-** Normalized fold change relative to control.

**(C)** Chromatin-bound ORC6 per cell in G1 from control or treated RPE1 cells expressing c-MYC at 20 ng/mL for 24 hours. The aggregate of two biological replicates was randomly downsampled to 4000 cells per sample.

**(D)** Immunoblotting for total ORC2 and ORC4 in lysates of control or doxycycline-treated RPE1 cells at 20 ng/mL for 24 hours. n=1.

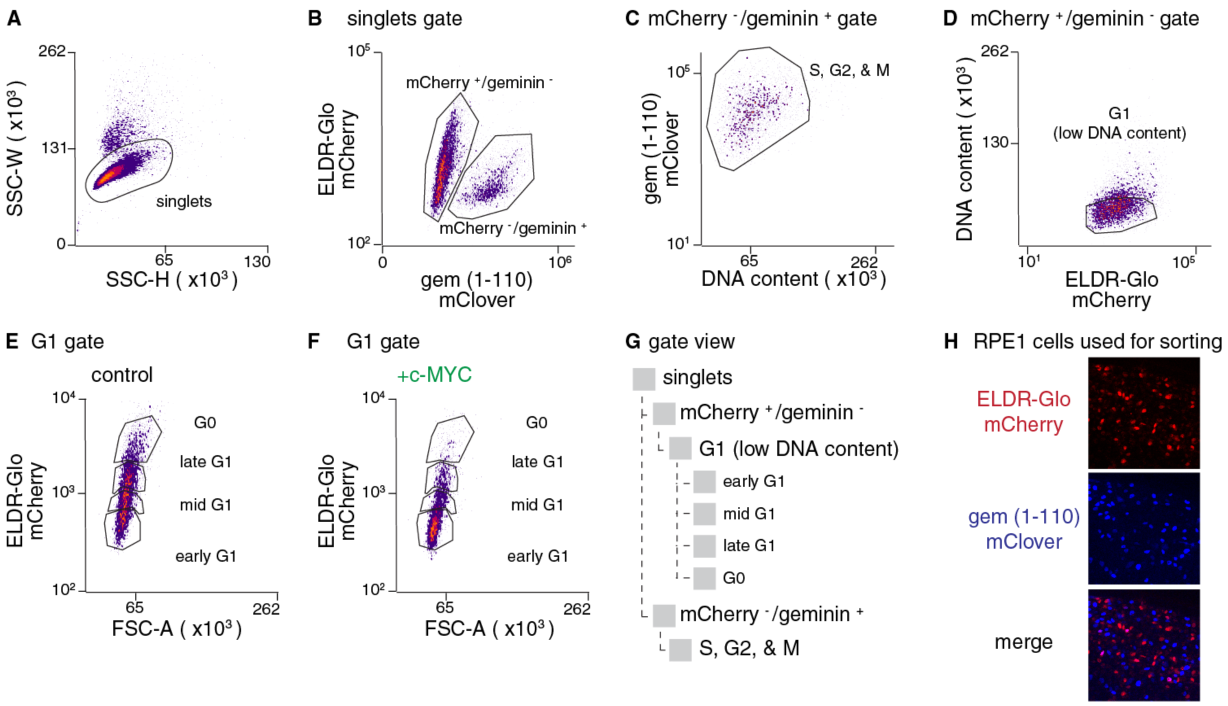


**Supplementary Fig. S8. Gating strategy for sorting cells in G1 subphases, G0, and S-M phases.**

**(A-G)** Fluorescence-activated cell sorting (FACS). Singlets were gated in (A) from all cells and doublets were excluded. Two gates were created in (B): mCherry positive/geminin-negative "mCherry ⁺/ geminin ⁻" cells which correspond to G0/G1 cells, mCherry-negative/geminin-positive "mCherry ⁻/ geminin ⁺" cells which correspond to S, G2 & M cells in (C) as illustrated in Fig. 2D. mClover is the color of the gem (1-110) biosensor. In (D), G1 cells were gated as the mCherry ⁺/ geminin ⁻ cells with low DNA content to further exclude any doublets. In (E) and (F), the early, mid, late G1 and G0 gates were placed arbitrarily but kept constant between the samples for control (E) or c-MYC overexpressing cells (F). The G0 gate was placed as the top ~15% of control cells with the highest mCherry intensity based on negative phosphorylated RB staining from Johnson *et al.*[30] signal. The gate view is shown in (G) for illustration. Representative of n>3.

**(H)** Image of RPE1 cells expressing the two mutually-exclusive biosensor signals: ELDR-Glo mCherry and the gem (1-110) mClover (depicted in blue).

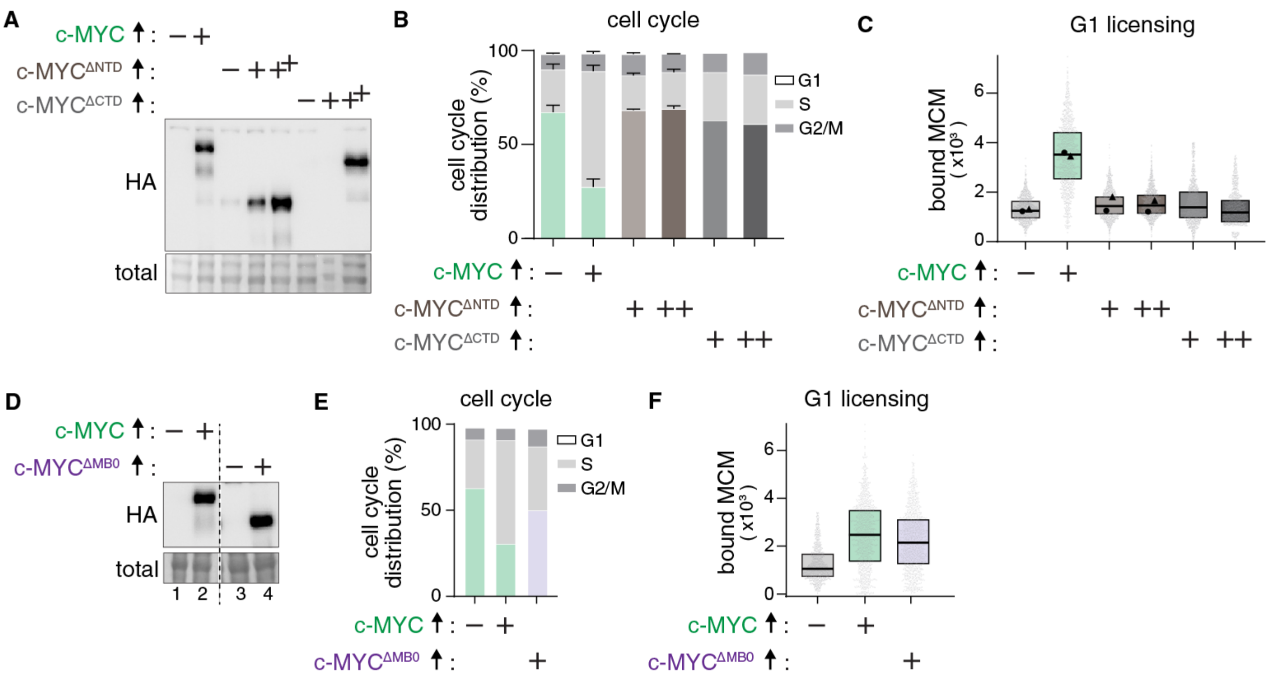


**Supplementary Fig. S9**

**The N- and C- terminal domains, but not the MB0, of c-MYC are essential for origin licensing acceleration.**

**(A)** Immunoblotting for HA to detect ectopic $^{2xHA}$c-MYC or $^{2xHA}$c-MYC harboring a deletion of the NTD ($^{2xHA}$c-MYC$^{\Delta NTD}$) or the CTD ($^{2xHA}$c-MYC$^{\Delta CTD}$) in control ("-") or doxycycline-treated RPE1 cells for 24 hours at 20 ng/mL ("+") for $^{2xHA}$c-MYC or at 20 ("+") or 100 ng/mL ("++") for $^{2xHA}$c-MYC$^{\Delta NTD}$ and $^{2xHA}$c-MYC$^{\Delta CTD}$.

**(B)** Cell cycle phase distribution from the same samples in (A); mean with error bars ± SEM.

**(C)** Chromatin-bound MCM per cell in G1 from the same samples in (B). Cells in each sample were randomly downsampled to 1000 or 2000 cells per sample for the c-MYC$^{\Delta CTD}$ or c-MYC$^{\Delta NTD}$ conditions, respectively. n=1 for c-MYC versus c-MYC$^{\Delta CTD}$. Data are combined from two biological replicates for c-MYC versus c-MYC$^{\Delta NTD}$.

**(D)** Immunoblotting for HA to detect ectopic $^{2xHA}$c-MYC or $^{2xHA}$c-MYC harboring a deletion of the MB0 domain ($^{2xHA}$c-MYC$^{\Delta MB0}$) in control or doxycycline-treated RPE1 cells at 20 ng/mL for 18 hours. Irrelevant lanes were manually removed from the blot image (dashed line).

**(E)** Cell cycle phase distribution from the same cells in (D). n=1.

**(F)** Chromatin-bound MCM per cell in G1 from the same samples in (E). Cells in each sample were randomly downsampled to 1000 cells per sample.

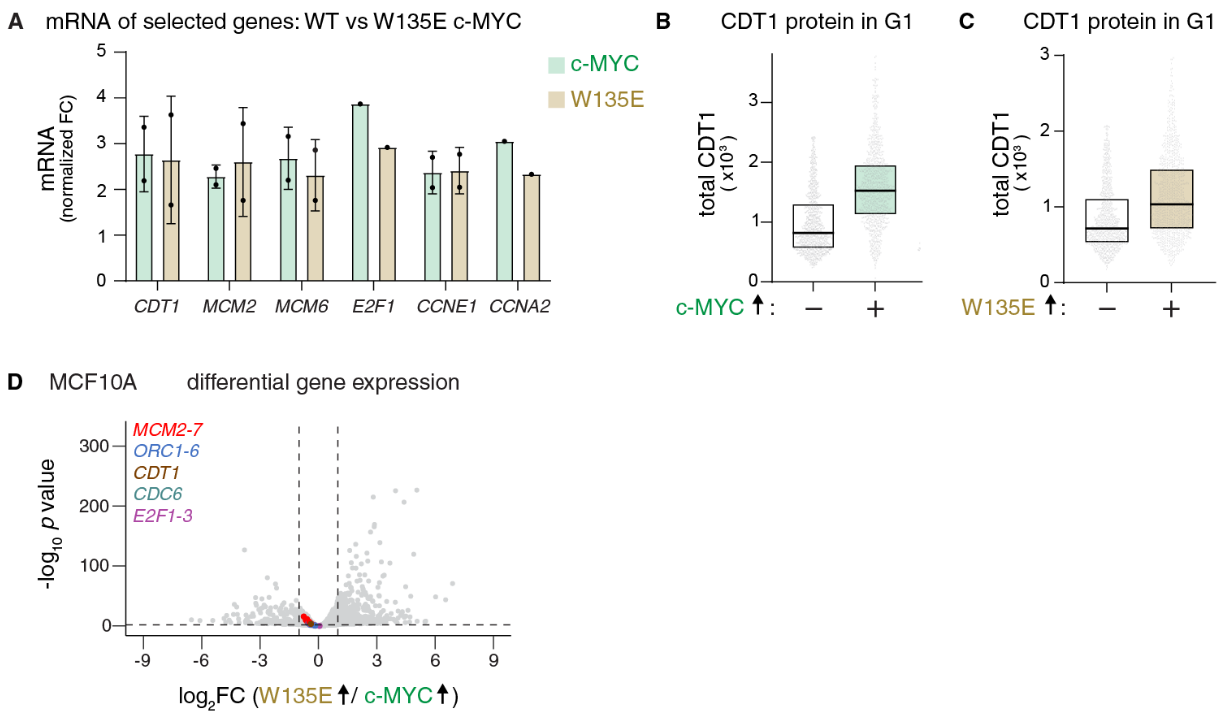


**Supplementary Fig. S10. Similar gene expression and CDT1 protein changes induced by WT and W135E c-MYC.**

**(A)** Bar graphs for the mRNA levels of the indicated genes in control or treated RPE1 cells expressing c-MYC or W135E at 20 ng/mL for 24 hours measured by qPCR. Transcript abundances in the treated samples were normalized to controls within each replicate set to 1 for relative fold change. Error bars display the mean ± SD of n=2 biological replicates, n=1 for *E2F1* and *CCNA2*.

**(B)** Total CDT1 protein levels per cell in G1 of control ("-") or RPE1 cells expressing c-MYC at 20 ng/mL for 24 hours ("+"). 1000 cells per sample. Fold-change in median CDT1 intensity relative to control: WT c-MYC = 1.77. n=1.

**(C)** Total CDT1 protein levels per cell in G1 of control or treated RPE1 cells expressing W135E at 20 ng/mL for 24 hours. 1000 cells per sample. Fold-change in median CDT1 intensity relative to control: W135E c-MYC = 1.41. n=1.

**(D)** Volcano plot showing the differentially expressed genes in MCF10A cells constitutively-expressing W135E or WT c-MYC from publicly-available data in Hinds *et al.* [26]. Origin licensing factors and E2F activator genes are color-coded. $Log_2$(fold change) threshold = ± 1, adjusted *p*-value threshold = 0.05.

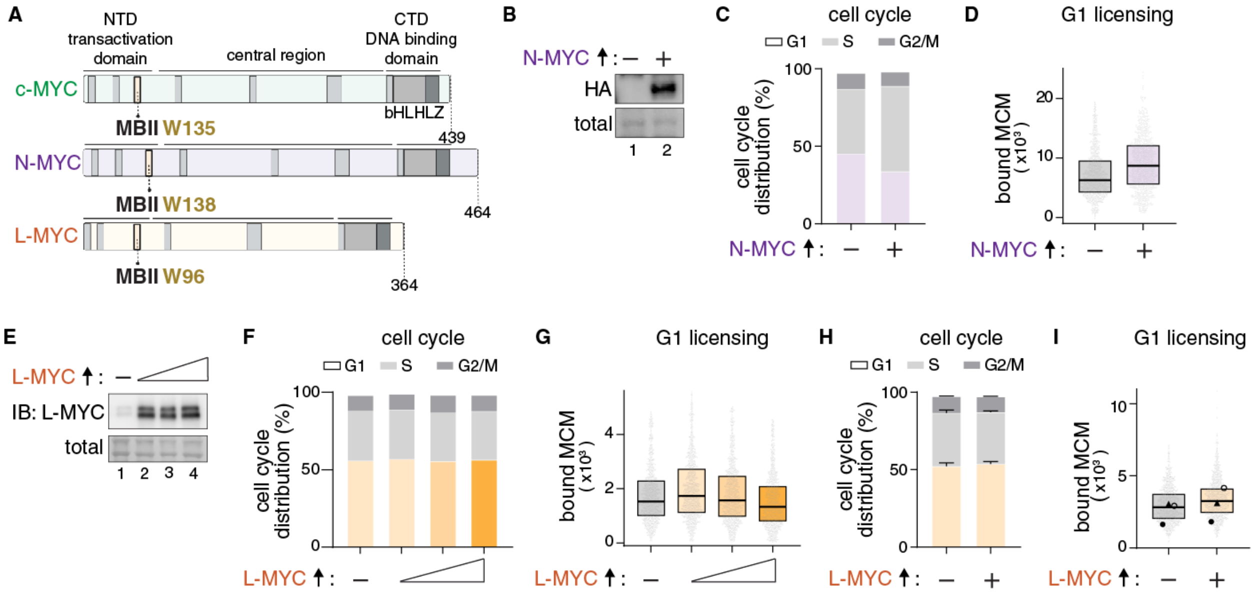


**Supplementary Fig. S11. c-MYC and N-MYC, but not L-MYC, expression accelerate licensing and shorten G1.**

**(A)** Schematic for c-MYC, N-MYC and L-MYC.

**(B)** Immunoblotting for HA to detect ectopic $^{2xHA}$-N-MYC in control or doxycycline-treated polyclonal RPE1 cells at 50 ng/mL for 19 hours.

**(C)** Cell cycle phase distribution of control or treated RPE1 cells overexpressing N-MYC at 100 ng/mL of doxycycline for 24 hours. n=1.

**(D)** Chromatin-bound MCM per cell in G1 from the same samples in (C). 1000 cells per sample. n=1.

**(E)** Immunoblotting for L-MYC to detect ectopic L-MYC in control (lane 1) or doxycycline-treated RPE1 cells at 20, 100, or 500 ng/mL (triangle) for 24 hours.

**(F)** Cell cycle phase distribution from the same samples in (E).

**(G)** Chromatin-bound MCM per cell in G1 from the same samples in (F). 1000 cells per sample. n=1.

**(H)** Cell cycle phase distribution of control or treated RPE1 cells overexpressing L-MYC at 20 ng/mL of doxycycline for 24 hours; mean with error bars ± SEM. n=3.

**(I)** Chromatin-bound MCM per cell in G1 from (H). The aggregate of three biological replicates was randomly downsampled to 3000 cells per sample. Statistical significance between control versus L-MYC expression was determined by a ratio paired *t*-test on the median fluorescence intensities from three independent biological replicates; $p$=0.21.

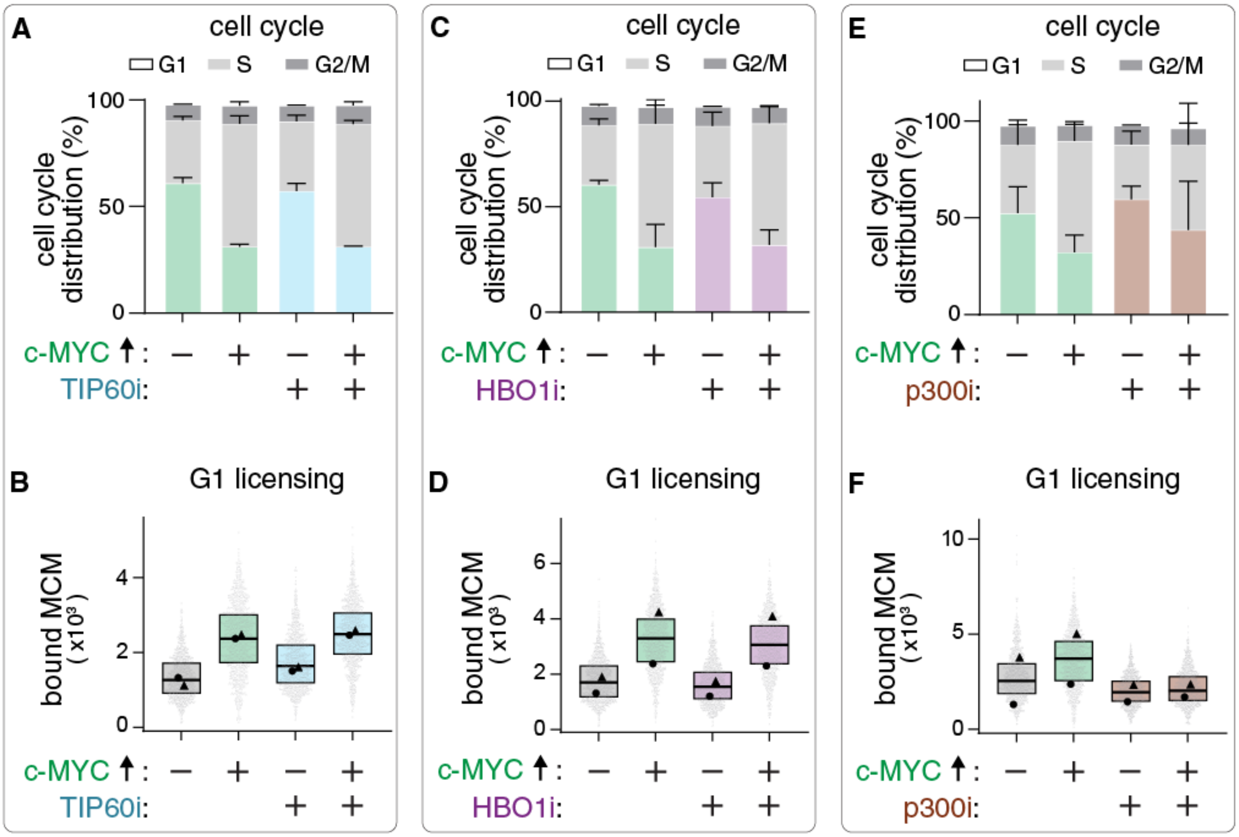


**Supplementary Fig. S12. The acetyltransferase p300, but not TIP60 or HBO1, is required for c-MYC to accelerate licensing.**

**(A)** Cell cycle phase distribution; mean with error bars ± SEM. RPE1 cells were treated with the TIP60 inhibitor (NU9056) at 25 µM, or 20 ng/mL doxycycline to express c-MYC, or both for 16-18 hours as indicated by "+". n=2.

**(B)** Chromatin-bound MCM per cell in G1 from the same samples in (A). 2000 cells per sample. Data are combined from two biological replicates; n=2.

**(C)** Cell cycle phase distribution; mean with error bars ± SEM. RPE1 cells were treated with the HBO1 inhibitor (WM-3835) at 5 µM, or 20 ng/mL doxycycline to express c-MYC, or both for 24 hours as indicated. n=2.

**(D)** Chromatin-bound MCM per cell in G1 from the same samples in (C). 2000 cells per sample. Data are combined from two biological replicates; n=2.

**(E)** Cell cycle phase distribution; mean with error bars ± SEM. RPE1 cells were treated with the p300 inhibitor (C646) at 10 µM, or 20 ng/mL doxycycline to express c-MYC, or both for 24 hours as indicated. n=2.

**(F)** Chromatin-bound MCM per cell in G1 from the same samples in (E). 2000 cells per sample. Data are combined from two biological replicates; n=2.

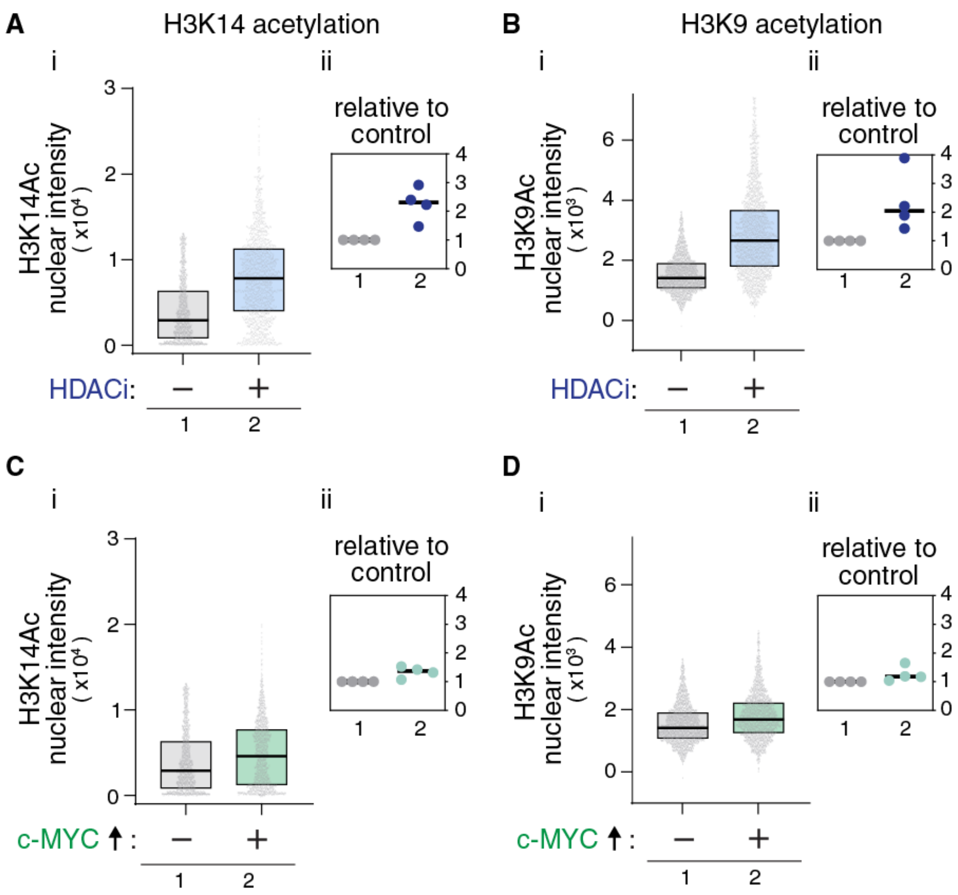


**Supplementary Fig. S13. c-MYC expression or histone deacetylase treatment increase cellular histone H3K14 and H3K9 acetylation.**

**(A) i-** Histone H3K14Ac levels in control and cells treated with the HDACi (TSA) at 300 nM for 4 hours. Representative of n=4. 1200 cells / sample. Ratio paired *t*-test on the median fluorescence intensities from four independent biological replicates; $p$=0.01. **ii-** Normalized fold change relative to control.

**(B)** As in (A), but measuring histone H3K9Ac levels. 2000 cells / sample. Ratio paired *t*-test on the median fluorescence intensities from four independent biological replicates; $p$=0.03.

**(C) i-** Histone H3K14Ac levels in control cells versus cells expressing c-MYC at 20 ng/mL for 24 hours. Representative of n=4. 1200 cells / sample. Ratio paired *t*-test on the median fluorescence intensities from four independent biological replicates; $p$=0.03. **ii-** Normalized fold change relative to control.

**(D)** As in (C), but measuring histone H3K9Ac levels. 2000 cells / sample. Ratio paired *t*-test on the median fluorescence intensities from four independent biological replicates; $p$=0.11. Note: the HDACi-treated and c-MYC-induced samples for each histone mark shown were run under the same experimental conditions on the same day.

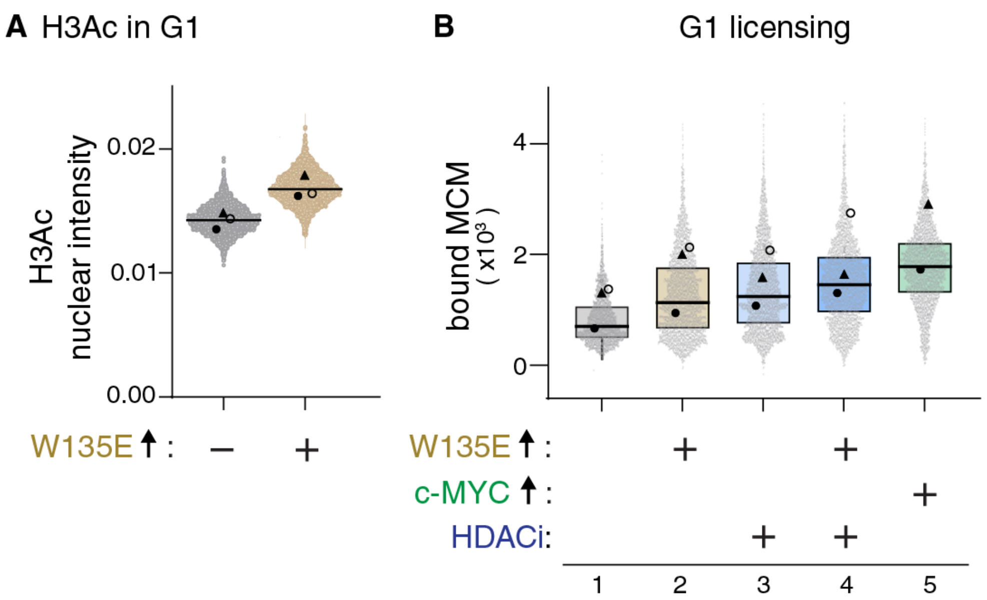


**Supplementary Fig. S14. W135E c-MYC is impaired for licensing acceleration despite H3Ac induction.**

**(A)** Histone H3Ac quantified by immunofluorescence in control or W135E-expressing G1 cells at 20 ng/mL of doxycycline for 16-24 hours (as in Fig. 4F). Data are the aggregate of three biological replicates, and the global median is marked; control=7015 nuclei, + W135E=5266 nuclei. Statistical significance between control versus W135E was determined by a ratio paired *t*-test on the median fluorescence intensities from three independent biological replicates; $p$=0.013.

**(B)** Chromatin-bound MCM per cell in G1 by analytical flow cytometry. RPE1 cells were treated with 20 ng/mL of doxycycline to express WT c-MYC, or W135E c-MYC for 24 hours, or treated with the HDACi (TSA) for 6 hours at 300 nM, washed off then released for 18 hours or treated with 20 ng/mL of doxycycline for 18 hours to overexpress W135E. Treatment combinations are as shown. The aggregate of three biological replicates was randomly downsampled to 4200 cells per sample for samples 1-4, and 3400 cells for sample 5. Statistical significance between W135E versus HDACi, or W135E versus W135E pre-treated with the HDACi was determined by a ratio paired *t*-test on the median fluorescence intensities from three independent biological replicates; $p$=0.73, $p$=0.52, respectively.

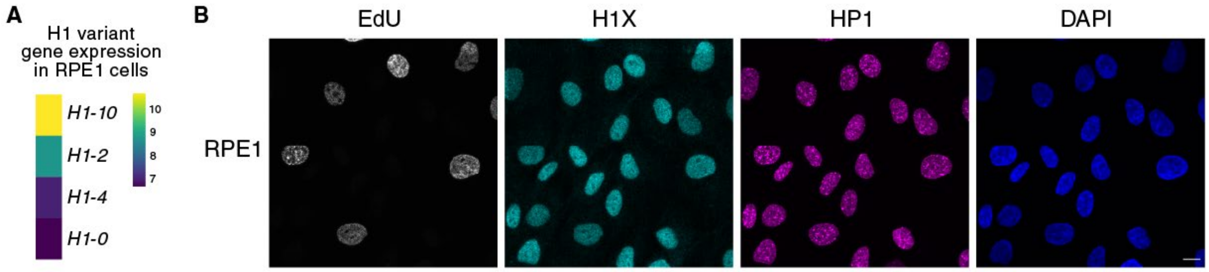


**Supplementary Fig. S15. H1X is the predominant H1 variant in RPE1 cells and displays uniform nuclear localization.**

**(A)** Normalized counts for histone H1 variant mRNAs in RPE1 control cells. RNAseq data from four biological replicates. The other somatic H1 variants *(H1-1, H1-3, and H1-5)* were below the detection definition (see Methods). *H1-10*: gene name for H1X.

**(B)** Representative immunofluorescence images for chromatin-bound H1X and HP1 (heterochromatin marker) in RPE1 cells. S-phase cells were identified by EdU signal. DNA were counterstained with DAPI. Scale bar, 10 μm. Representative of 3 biological replicates.

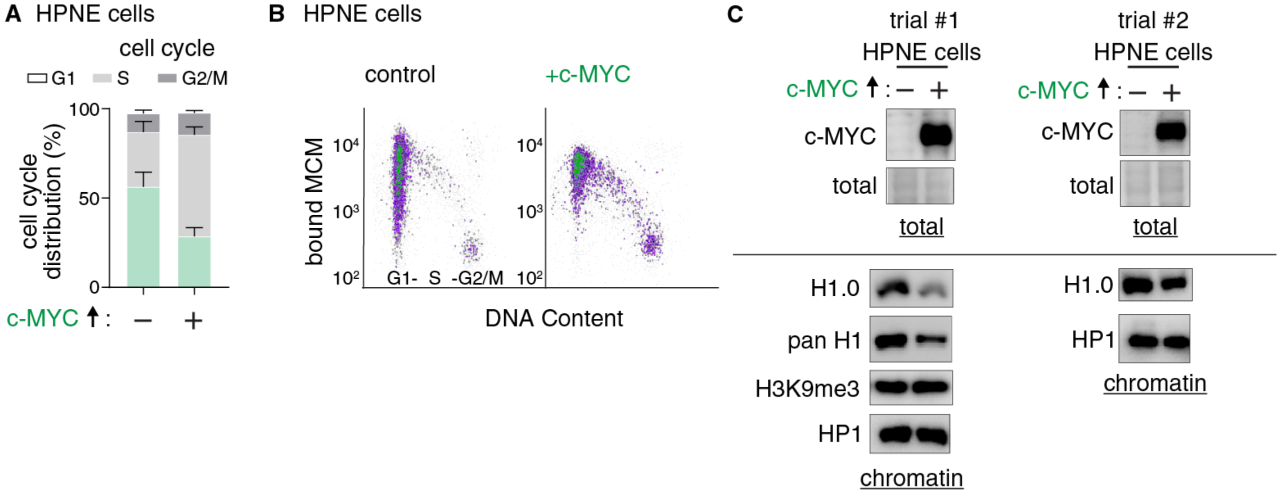


**Supplementary Fig. S16. c-MYC expression in HPNE cells shortens G1, evicts H1.0 from chromatin, and accelerates licensing.**

**(A)** Cell cycle phase distribution of control or treated HPNE cells with 1 µg/mL of doxycycline for 24 hours to express c-MYC; mean with error bars ± SEM. n=2.

**(B)** Representative chromatin-bound MCM per cell for cells treated as in (A). Cells were stained for bound MCM2 and DAPI for DNA content. Raw data are shown as a representative of two biological replicates. Each plot has 10,000 cells.

**(C)** Whole cell lysates or chromatin fractions were analyzed by immunoblotting for the indicated proteins in control or doxycycline-treated HPNE cells at 1 µg/mL for 24 hours. Ponceau S was used to detect total protein content for the whole cell lysates. HP1 was used as a loading control for the chromatin fractions. Two independent biological replicates are shown.

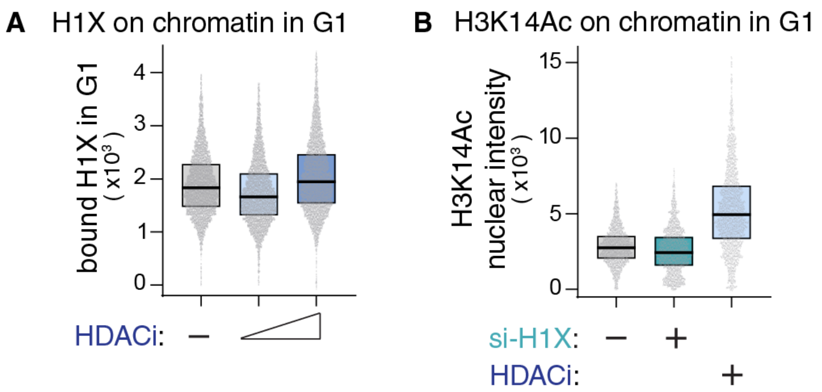


**Supplementary Fig. S17. Histone hyperacetylation and histone H1 eviction are independent.**

**(A)** Chromatin-bound H1X per cell in G1 of control or treated RPE1 cells with the histone deacetylase inhibitor HDACi (TSA) at 300 nM or 1 µM for 6 hours. 5000 cells per sample. n=1.

**(B)** Chromatin-bound H3K14Ac levels in G1 from cells treated with control or H1X siRNAs at 100 nM for 48 hours. Cells treated with trichostatin A at 300 nM for 3 hours are shown as a positive control. 2000 cells per sample. n=1

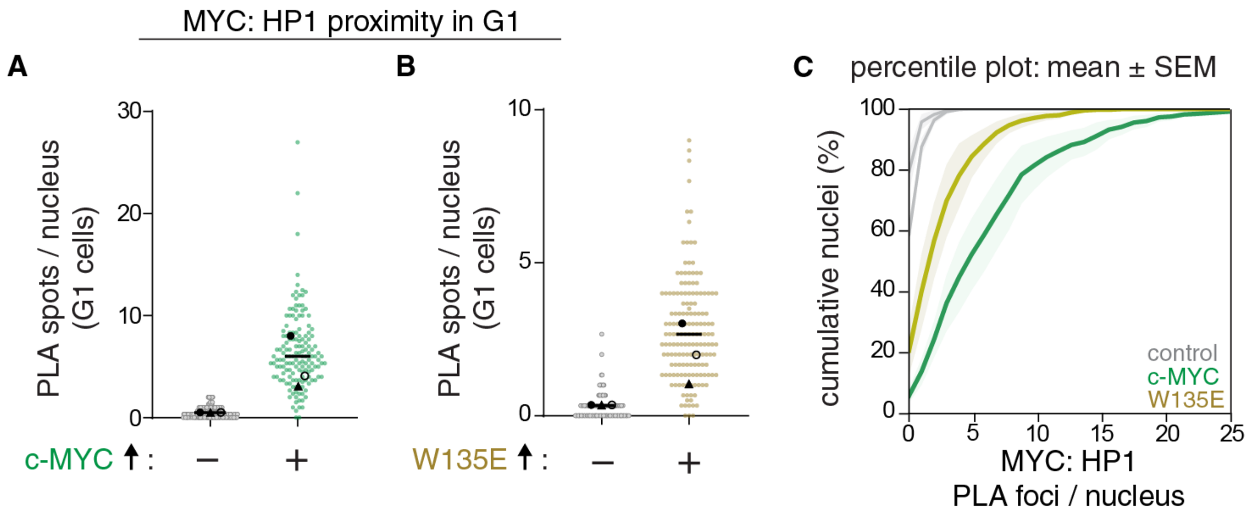


**Supplementary Fig. S18. Overexpressed WT and W135E c-MYC localize to heterochromatin.**

**(A)** WT c-MYC-HP1 proximity ligation assay (PLA) foci per nucleus in G1 cells of control and WT c-MYC overexpressing cells (470 cells for control, 397 cells for + c-MYC). Line at median is shown in black. Data are combined from three independent biological replicates.

**(B)** As in (A) but comparing control to W135E c-MYC overexpressing cells. (397 cells for control, 443 cells for + W135E). Line at median is shown in black. Data are combined from three independent biological replicates. *t*-test of mean intensities;

**(C)** Replicate-aware percentile or Empirical Cumulative Distribution Function (ECDF) plot for the cumulative % of nuclei (y-axis) versus the number of PLA foci per nucleus (x-axis) for each condition in (A) and (B). Shaded areas represent the SEM across three biological replicates.
